# Antidepressant treatment modulates the gut microbiome and metabolome during pregnancy and lactation in rats with a depressive-like phenotype

**DOI:** 10.1101/501742

**Authors:** Anouschka S Ramsteijn, Eldin Jašarević, Danielle J Houwing, Tracy L Bale, Jocelien DA Olivier

## Abstract

**Background:** Up to 10% of women use selective serotonin reuptake inhibitor (SSRI) antidepressants during and after pregnancy to manage mood disorders, with possible implications for the developing offspring. The microbiota within the gastrointestinal tract contributes to the regulation of serotonin synthesis. However, the interaction between maternal depression, SSRI use, bacterial community composition, and availability of microbiota-derived metabolites during pregnancy and lactation is not clear and may be consequential to the long-term health of mother and offspring. To determine the impact of SSRI treatment on maternal microbial community dynamics, we conducted these studies in a rat model of maternal vulnerability (MV). All MV females are on a background of genetic vulnerability, where rats exposed to early life stress (sMV) develop a depressive-like phenotype. In adulthood, sMV- and control (cMV) females were treated with either the SSRI fluoxetine (FLX) or the vehicle (Veh) throughout pregnancy and lactation. High-resolution 16S ribosomal RNA gene sequencing and targeted metabolomic analysis were used to assess the fecal microbiome and metabolite availability, respectively.

**Results:** The diversity, structure, and composition of the fecal bacterial community differed between pregnancy and lactation. Shifts in microbiota composition were accompanied by changes in fecal metabolite availability. FLX altered some key features of the transition from pregnancy to lactation, but only in females exposed to early life stress (sMV-Veh vs sMV-FLX). For instance, sMV-FLX females had lower fecal availability of the amino acids serine, proline, and aspartic acid than sMV-Veh females. These metabolite concentrations correlated negatively with the relative abundance of bacterial taxa *Prevotella, Ruminococcus*, and *Oscillospira*.

**Conclusions:** Our studies demonstrate an important relationship between antidepressant use during the perinatal period and maternal fecal metabolite availability in sMV rats, possibly through parallel changes in the maternal gut microbiome. Since maternal microbial metabolites contribute to health outcomes in offspring, insults to the maternal microbiome by SSRIs might have inter-generational consequences.

## Background

Dysregulation of the neurotransmitter serotonin in the brain is a widely recognized hallmark of major depressive disorder [1, 2]. Selective serotonin reuptake inhibitor (SSRI) antidepressants target the serotonin transporter (SERT), thereby influencing the serotonergic tone [3]. Serotonin (also referred to as 5-hydroxytryptamine or 5-HT) and SERT are particularly well-known for playing crucial regulatory roles in the brain, but they also act in many peripheral tissues. Nearly 95% of the body’s serotonin resides within the gastrointestinal tract, where it is mainly produced by enterochromaffin cells [4]. Interestingly, germ-free and antibiotic-treated rodents show lower levels of colonic and serum serotonin. This is reversible by microbial colonization, suggesting an important role for specific members of the gut microbiome in modulating serotonin availability [5–11]. Indeed, microbiota synthesize tryptophan, the precursor of serotonin, and promote serotonin biosynthesis in the host organism [8, 9]. Conversely, host serotonin signaling impacts the gut microbiota composition, as we have shown using rats with reduced or no SERT gene expression [12]. Several studies have found alterations in the gut microbiota composition of patients with major depression versus healthy controls [13, 14]. The same has been observed in rodents with depressive-like symptoms [15]. Moreover, a recent *in vivo* study showed that SSRIs modulate the gut microbiota composition, an effect that may be related to *in vitro* antimicrobial properties of SSRIs [16–20].

The maternal gut microbiota synthesizes a variety of metabolites that reach the systemic circulation during pregnancy and lactation, and influence offspring innate immune development [21, 22]. Environmental and pharmacological insults disrupt the compositional and functional states of the maternal gut microbiota, as recently demonstrated by several animal studies focusing on stress, diet, and antibiotic use during pregnancy [23–27]. Despite clear evidence that the microbiome interacts with serotonin homeostasis, it is currently unknown whether alterations in serotonin homeostasis, such as those often co-occurring with depression or resulting from SSRI use, affect the maternal gut microbiome during the perinatal period [5, 8–12]. An estimated 7-13% of women suffer from a major depressive disorder in the perinatal period and 1-10% of pregnant women take SSRI antidepressants, further highlighting the necessity to better understand the effect of SSRI medication during this period on the gut microbiota [28–31].

We aimed to investigate whether a depressive-like phenotype, SSRI antidepressant treatment, and their combination affect the microbial community composition and function during pregnancy and the postpartum period. To this end, we used a rat model of maternal vulnerability (MV) in combination with early life stress by maternal separation (sMV) or control handling (cMV). In adulthood, sMV females show a lower sucrose preference and lower neural growth factor gene expression in the basolateral amygdala and paraventricular nucleus of the brain than cMV females [32]. Thus, sMV females show anhedonia, which is an endophenotype associated with depressive symptoms. We use sMV rats as a model for depression, allowing us to study the separate effects of the maternal depressive-like phenotype and SSRI use during pregnancy, as well as the combination of both.

In the current study, adult female sMV and cMV rats were treated daily with fluoxetine (FLX), a commonly used SSRI, or vehicle (Veh) throughout pregnancy and lactation. There were 4 groups of MV females; the control group that was treated with vehicle (cMV-Veh), the maternally separated group that was treated with vehicle (sMV-Veh), the control group that was treated with fluoxetine (cMV-FLX), and the maternally separated group that was treated with fluoxetine (sMV-FLX). Weekly fecal samples were collected for 16S rRNA gene sequencing. In addition, targeted metabolomic analysis of amino acids, short-chain fatty acids and bile acids was done on a subset of samples. We examined the hypothesis that depressive-like symptoms and antidepressant treatment during pregnancy and lactation affect the maternal fecal microbiome and its functional capacity. Specifically, we hypothesized that (1) pregnancy and lactation have distinct fecal microbial signatures; (2) early life stress and FLX alter the fecal microbial signatures of pregnancy and lactation, and the combination of early life stress and FLX has the most pronounced effect; (3) pregnancy and lactation have distinct fecal metabolic signatures; (4) early life stress and FLX treatment alter the fecal metabolic signatures of pregnancy and lactation, and the combination of early life stress and FLX has the most pronounced effect; and (5) the abundances of microbial taxa correlate with the concentrations of fecal metabolites.

## Methods

### Experimental animals

The animals came from our colony of serotonin transporter knockout (SERT^-/-^, Slc6a41^Hubr^) Wistar rats at the University of Groningen (Groningen, the Netherlands), and were derived from outcrossing SERT^+/-^ rats [33]. Animals were supplied *ad libitum* with standard lab chow (RMH-B, AB Diets; Woerden, the Netherlands) and water and were kept on a 12:12 h light-dark cycle (lights off at 11:00 h), with an ambient temperature of 21 ± 2 °C and humidity of 50 ± 5%. Cages were cleaned weekly, and animals were provided with a wooden stick for gnawing (10−2−2cm) and nesting material (Enviro-dri™, Shepherd Specialty Papers, Richland, MI, USA). During pregnancy, females were housed individually in type III Makrolon cages (38.2−22.0−15.0cm) and stayed with their pups until postnatal day (PND)21. Animals were weaned at PND21 and group housed in same-sex cages of 3-5 animals in type IV (55.6−33.4−19.5cm) Makrolon cages. Genotyping of the animals was performed as described previously [12]. All experimental procedures were approved by the Institutional Animal Care and Use Committee of The University of Groningen and were conducted in agreement with the Law on Animal Experiments of The Netherlands.

### Early life stress by maternal separation in a rat model of maternal vulnerability

Our rat model of maternal vulnerability (MV) consists of heterozygous SERT knockout (SERT^+/-^) female rats. MV rats show 40-50% lower SERT gene expression levels than wildtypes [34]. These levels are similar to those of a considerable portion of the human population, who carry at least one short allele of the SERT gene [35]. When human carriers of this allele experience severe stressful life events, they are at higher risk of developing symptoms of depression [36]. We mimic this gene-by-environment interaction contributing to depression in MV rats by exposing them to early life stress [37]. The early life stress protocol was conducted as previously described [12]. In short, SERT^+/-^ females were mated with SERT^+/-^ males (F0). Females were left undisturbed until delivery, which was defined as PND0. From PND2 to PND15, pups were either separated from the dam for 6 hours per day, or handled 15 minutes per day (Figure 1a). The SERT^+/-^ female pups from these nests (F1) then matured to become the sMV- and cMV females.

**Figure 1:**
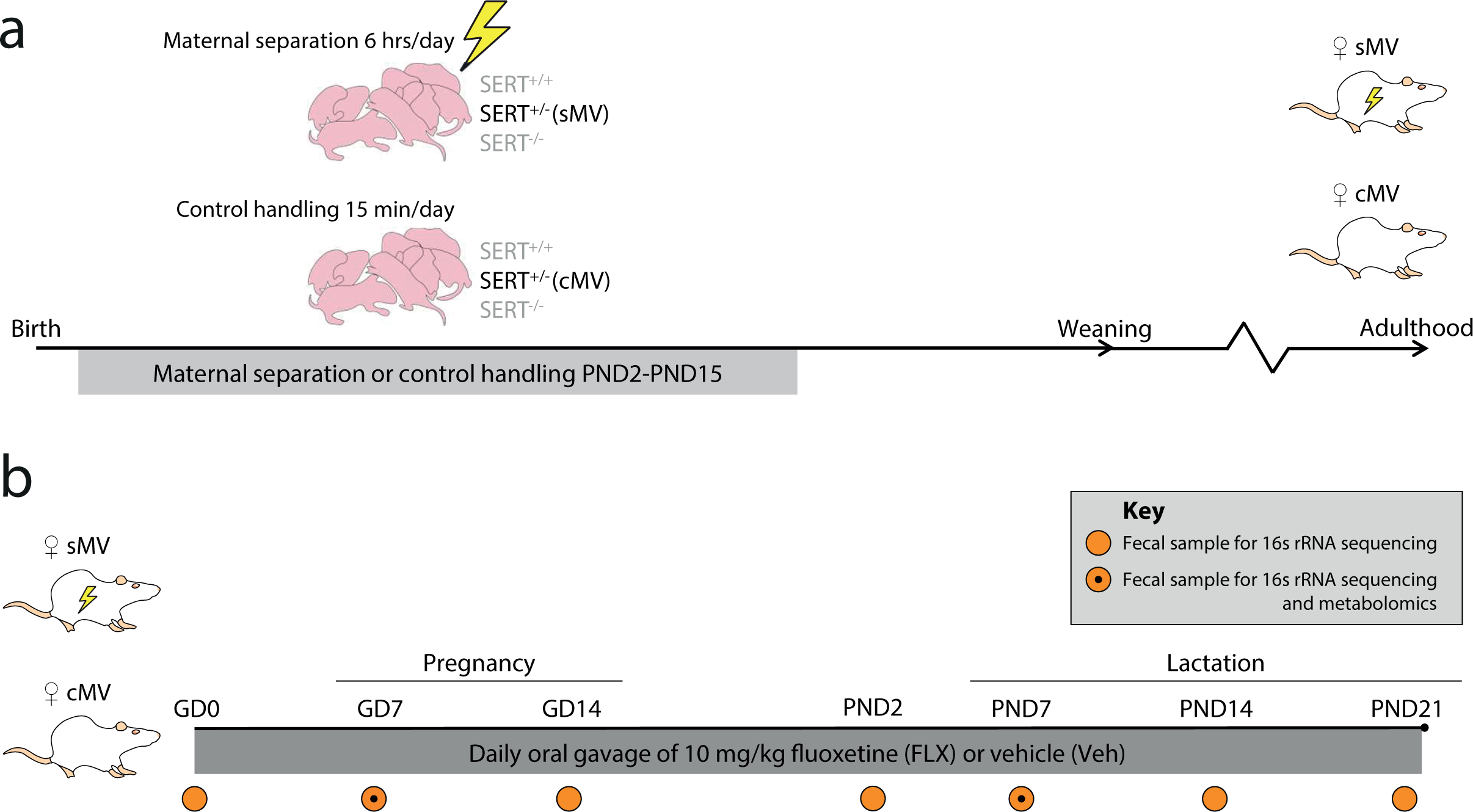
Overview of study design and sampling schedule. (a) Early life stress protocol. SERT^+/-^ females were crossed with SERT^+/-^ males, yielding nests with offspring genotypes SERT^+/+^, SERT^+/-^, and SERT^-/-^. The SERT^+/-^ females are our model of maternal vulnerability (MV). From postnatal day (PND) 2 to PND15, pups were either maternally separated for 6 hours per day (early life stress - sMV) or control handled (cMV) for 15 minutes per day. Pups were weaned at PND21. sMV- and cMV females were group housed (same treatment) until adulthood. **(b) Fluoxetine treatment during pregnancy and lactation, and fecal sampling schedule.** Adult sMV and cMV females (N=14-18/group) were crossed with wildtype males. Through pregnancy and lactation, from gestational day (GD)1 until PND21, females received a daily oral injection of either 10 mg/kg fluoxetine (FLX) or methylcellulose (MC, Veh). Thus, there were 4 groups of females: cMV-Veh, sMV-Veh, cMV-FLX, and sMV-FLX (N=6-11/ group). Fecal pellets for 16s rRNA gene sequencing were freshly collected before conception at GD0, during pregnancy at GD7 and GD14, at PND2, and during lactation at PND7, PND14 and PND21 (N=192 in total). Selected fecal samples from GD7 and PND7 were also used for metabolomic analysis (N=4-5/group per time point, N=36 in total).

### Breeding and fluoxetine treatment

sMV and cMV females were mated with wildtype males. Females were between the age of 3 and 6 months, and mated when in estrus (checked with a disinfected impedance meter, model MK-11, Muromachi, Tokyo, Japan). This was termed gestational day (GD)0. The males were removed after 24 hours, and the females stayed isolated in a type III cage. Throughout pregnancy and lactation, from GD1 until PND21, the dams were weighed and received an oral gavage of either 10 mg/kg fluoxetine (FLX, Fluoxetine 20 PCH, Pharmachemie BV, the Netherlands) or vehicle daily at 11:00AM (Figure 1b). Methylcellulose (MC; Sigma Aldrich Chemie BV, Zwijndrecht, the Netherlands) was used for vehicle injections since it is the constituent of the fluoxetine capsule. FLX (5 mg/mL) and MC (1%) solutions were prepared with autoclaved water. The gavage volume ranged from 0.9 mL to 2.0 mL (depending on body weight). Animals were treated orally by gently picking up the animal without restraint, and using flexible PVC feeding tubes (40 cm length, Vygon, Valkenswaard, the Netherlands) in order to minimize discomfort and stress in the animals. In total, 4 groups of MV females were sampled for this study: cMV-Veh (N=20), sMV-Veh (N=13), cMV-FLX (N=34) and sMV-FLX (N=25). However, the final number of animals selected for 16S rRNA gene sequencing for this study were 11 cMV-Veh, 8 sMV-Veh, 7 cMV-FLX and 6 sMV-FLX. Because littermates can share microbiotas, no females from the same litter were used within each group.

### Fecal sample collection

Fecal samples to be sequenced were collected at GD0 (before conception), GD7, GD14, PND2, PND7, PND14 and PND21 (Figure 1b). We chose not to collect samples closer to the day of giving birth to not induce unnecessary stress. The samples from GD7 and GD14 were grouped as pregnancy and the samples from PND7, PND14 and PND21 as lactation, since we observed no significant differences within these periods in terms of alpha diversity, beta diversity, and composition of the microbiome (data not shown), as was also shown in a previous human study [38]. Conversely, the samples from PND2 were left out of the analysis, since they did not fit either category (data not shown). The samples were freshly collected directly from the animal between 10:00AM and 11:00AM, and precautions were taken to minimize sample contamination, such as the use of gloves and disinfection of work surfaces. The samples were placed in clean 2.0 mL Safe-Lock Eppendorf tubes (Nijmegen, The Netherlands), snap frozen in liquid nitrogen, and stored at −80 °C.

### 16S rRNA gene sequencing and analysis

Genomic DNA was isolated from fecal pellets using the PSP® Spin Stool DNA Kit (STRATEC Molecular GmbH, Berlin, Germany), according to the manufacturer’s instructions for difficult to lyse bacteria. About 100-200 mg fecal matter per sample was used for the DNA isolation; the remainder of the sample was stored for metabolite measurements. A dual-index sequencing strategy targeting the V4 region of the 16S rRNA gene was employed, using barcoded primers for the Illumina platform, as described previously [39]. Library concentration was quantified using Qubit and an additional primer dimer clean-up step was conducted using the AMPure XP beads according to the manufacturer’s protocol (Beckman Coulter, Brea, CA, USA). Sequencing was executed on a MiSeq instrument (Illumina, San Diego, CA) using 250 base paired-end chemistry at the University of Pennsylvania Next Generation Sequencing Core. For quality control purposes, a sample of the Human Microbiome Project Mock Community was included as a positive control, and water as a negative control. Quality filtering and chimera checking yielded 7,867,454 quality-filtered sequences with a mean ± SD depth of ± 2560 reads per sample. 16S rRNA gene analysis was performed using mothur [40] and QIIME version 1.9.1 (Quantitative Insights Into Microbial Ecology [41]) as described previously [23]. In short, Operational Taxonomic Units (OTUs) were defined with 97% sequence similarity using clustering method CD-HIT. The samples were rarified to 1,000 sequences per sample before calculating diversity metrics. The Shannon diversity index was used to calculate alpha diversity, and weighted UniFrac distances were used to calculate beta diversity.

### Statistical analysis

To identify the fecal microbial OTUs that differentiate between pregnancy and lactation, the machine-based learning algorithm Random Forests was used in R version 3.4.1 as previously described [23, 42–44]. OTU importance was ranked by the percent increase in prediction error of the model as a result of removal of that particular OTU from the model. The resulting model, derived from cMV-Veh samples, was then applied to OTU tables from the other groups, to see how early life stress, FLX and their combination alter the microbial signature of pregnancy and lactation. The R package “gplots” was used to plot OTU relative abundances as heat maps [45]. To predict the metabolic capacity of the gut microbiome during pregnancy and lactation, a computational approach known as Phylogenetic Investigation of Communities by Reconstruction of Unobserved States (PICRUSt [46]) was used. This algorithm uses the 16S rRNA data to predict the total metagenomic content, and these metagenomes are then mapped onto functional Kyoto Encyclopedia of Genes and Genomes (KEGG) pathways in order to predict the full functional capacity of the microbial community, as previously described [23]. Random Forests were applied to the dataset of predicted metabolic pathways to identify those that shift from pregnancy to lactation in the cMV-Veh group. Then, the impact of early life stress, FLX, and their combination on the proportional representation of these predicted microbial metabolic pathways was plotted in a heat map.

### Targeted metabolomics

To draw associations between bacterial community composition, predicted gut microbial metabolic capacity, and gut metabolite output, fecal metabolites were quantified in a subset of samples. GD7 was chosen as a representative time point for pregnancy, and PND7 for lactation. Fecal samples weighing about 300 mg from 5 randomly selected animals per treatment group were used (N=36 in total). Targeted metabolomics was performed by the Microbial Culture and Metabolomics Core as part of the PennCHOP Microbiome Program at the University of Pennsylvania via ultra-performance liquid chromatography (Acquity UPLC system, Waters Corporation, Milford, MA). Amino acid concentrations were quantified with an AccQ-Tag Ultra C18 1.7 μm 2.1−100mm column and a photodiode array detector. Analysis was performed using the UPLC AAA H-Class Application Kit (Waters Corporation, Milford, MA) according to manufacturer’s instructions. The limit of detection was 1 nmol/g stool. Bile acid concentrations were measured using a Cortecs UPLC C-18+ 1.6 mm 2.1 x 50 mm column, a QDa single quadrupole mass detector and an autosampler (192 sample capacity). The flow rate was 0.8 mL/min, the injection volume was 4 μL, the column temperature was 30°C, the sample temperature was 4°C, and the run time was 4 min per sample. Eluent A was 0.1% formic acid in water, eluent B was 0.1% formic acid in acetonitrile, the weak needle wash was 0.1% formic acid in water, the strong needle wash was 0.1% formic acid in acetonitrile, and the seal wash was 10% acetonitrile in water. The gradient was 70% eluent A for 2.5 minutes, gradient to 100% eluent B for 0.6 minutes, and then 70% eluent A for 0.9 minutes. The mass detection channels were: +357.35 for chenodeoxycholic acid and deoxycholic acid; +359.25 for lithocholic acid; +385.38 for obeticholic acid; +514.37 for glycoobeticholic acid; +528.39 for tauroobeticholic acid; −407.5 for cholic, alphamuricholic, betamuricholic, gamma muricholic, and omegamuricholic acids; −432.5 for glycolithocholic acid; −448.5 for glycochenodeoxycholic and glycodeoxycholic acids; −464.5 for glycocholic acid; −482.5 for taurolithocholic acid; −498.5 for taurochenodeoxycholic and taurodeoxycholic acids; and −514.4 for taurocholic acid. Samples were quantified against standard curves of at least five points run in triplicate. The range of the assay was at least 50 nM – 10,000 nM; the limit of detection was < 50 nM, and; the limit of quantitation was >10,000 nM. The limit of detection was 0.5 nmol/g stool. Finally, short-chain fatty acids were quantified using a HSS T3 1.8 μm 2.1−150 mm column with a photodiode array detector and an autosampler (192 sample capacity). The flow rate was 0.25 mL/min, the injection volume was 5 μL, the column temperature was 40°C, the sample temperature was 4°C, and the run time was 25 min per sample. Eluent A was 100 mM sodium phosphate monobasic, pH 2.5; eluent B was methanol; the weak needle wash was 0.1% formic acid in water; the strong needle wash was 0.1% formic acid in acetonitrile; the seal wash was 10% acetonitrile in water. The gradient was 100% eluent A for 5 min, gradient to 70% eluent B from 5-22 min, and then 100% eluent A for 3 min. The photodiode array was set to read absorbance at 215 nm with 4.8 nm resolution. Samples were quantified against standard curves of at least five points run in triplicate. Concentrations in the samples were calculated as the measured concentration minus the internal standard. The limit of detection was 1 nmol/g stool.

### Analysis of metabolomics data

All metabolites that were detected in at least 40% of samples were included in the analysis. For these metabolites, missing values were replaced by half the minimum positive value found for that metabolite, assuming that missing values were the result of the concentration being below the detection limit. Random Forests were applied as described above to the metabolite concentration table from the cMV-Veh samples in order to reveal which individual metabolites were overrepresented during pregnancy or during lactation. The resulting model was then applied to the data from the treatment groups, and heat maps were generated. For predictive analysis, the open-access, online platform MetaboAnalyst 3.0 was used for pathway analysis based on the metabolomics data [47]. Metabolite Set Enrichment Analysis, a metabolomic version of Gene Set Enrichment Analysis, was used to investigate which sets of functionally related metabolites differ between pregnancy and lactation per treatment group [48].

### Additional statistical methods

Microbial alpha diversity was analyzed using a two-way ANOVA with period (pregnancy vs lactation) and treatment group as between subject factors. Uncorrected Fisher’s LSD was used for group comparisons within periods (cMV-Veh vs sMV-Veh, cMV-Veh vs cMV-FLX, cMV-Veh vs sMV-FLX, sMV-Veh vs sMV-FLX, cMV-FLX vs sMV-FLX within pregnancy and within lactation). Permutational multivariate analysis of variance using distance matrices (PERMANOVA) was used to analyze effects of pregnancy vs lactation and treatment group on weighted UniFrac distances [49]. PERMANOVA p-values were based on 999 Monte Carlo simulations. Multiple comparisons of OTU relative abundances and metabolite concentrations within pregnancy and within lactation between treatment groups were done by nonparametric Kruskal-Wallis tests followed by uncorrected Dunn’s tests (cMV-Veh vs sMV-Veh, cMV-Veh vs cMV-FLX, cMV-Veh vs sMV-FLX, sMV-Veh vs sMV-FLX, cMV-FLX vs sMV-FLX within pregnancy and within lactation). All of the OTUs and metabolites analyzed in this way differed between pregnancy and lactation in the cMV-Veh group, following from the Random Forests analysis; further statistics between periods were not conducted. A t-test was used to analyze the GD0 alpha diversity difference between cMV and sMV females. For the abovementioned tests, GraphPad Prism version 7 (GraphPad Software, Inc., San Diego, CA) was used. The statistical significance is indicated as follows: \**p*<0.05; ** *p*<0.01; and *** *p*<0.001, **** *p*<0.0001. Data are presented as mean ± SEM, except OTU relative abundance which is plotted as median ± IQR.

## Results

We examined the hypothesis that depressive-like symptoms and antidepressant treatment during pregnancy and lactation affect the maternal fecal microbiome composition and function. Maternal vulnerability (MV) rats were used, since early life stress produces depressive-like symptoms and shifts in fecal microbiome community composition in these animals [12, 32]. In short, MV females exposed to either maternal separation or control handling for the first 2 weeks of life were treated with fluoxetine or vehicle daily throughout gestation and the postpartum period (Figure 1a,b). Thus, there were four groups of MV females: control (cMV-Veh), maternally separated and vehicle-treated (sMS-Veh), fluoxetine-treated (cMV-FLX) and maternally separated and fluoxetine-treated (sMV-FLX). It should be noted that we have been confronted with a number of unexpected and thus far unexplained deaths of some MV females as a result of fluoxetine treatment. About 25% of FLX-treated females died, even though there was no accumulation of fluoxetine or norfluoxetine levels in the blood over the course of treatment (data not shown). However, the individual females used for the current study did not show any abnormal response to FLX. Fecal pellets for 16s rRNA sequencing and analysis were collected during pregnancy at gestational day (GD)7, GD14, and during lactation at postnatal day (PND)7, PND14 and PND21. A subset of samples was used for targeted metabolomic analysis (Figure 1b).

### Pregnancy and lactation have distinct fecal microbial signatures

To examine whether pregnancy and lactation have distinct fecal microbial signatures, we first analyzed the 16S rRNA sequencing data of pregnancy versus lactation in terms of microbial alpha diversity and community structure over all 4 groups. Alpha diversity was higher during pregnancy than during lactation, as shown by the Shannon diversity index (Figure 2a, *p*<0.0001). The differences in phylogenetic structure between samples were explored by assessing weighted UniFrac distances. A separation between samples obtained during pregnancy versus lactation was revealed by Principle Coordinates Analysis (PCoA) (Figure 2b, PERMANOVA *p*=0.001).

**Figure 2:**
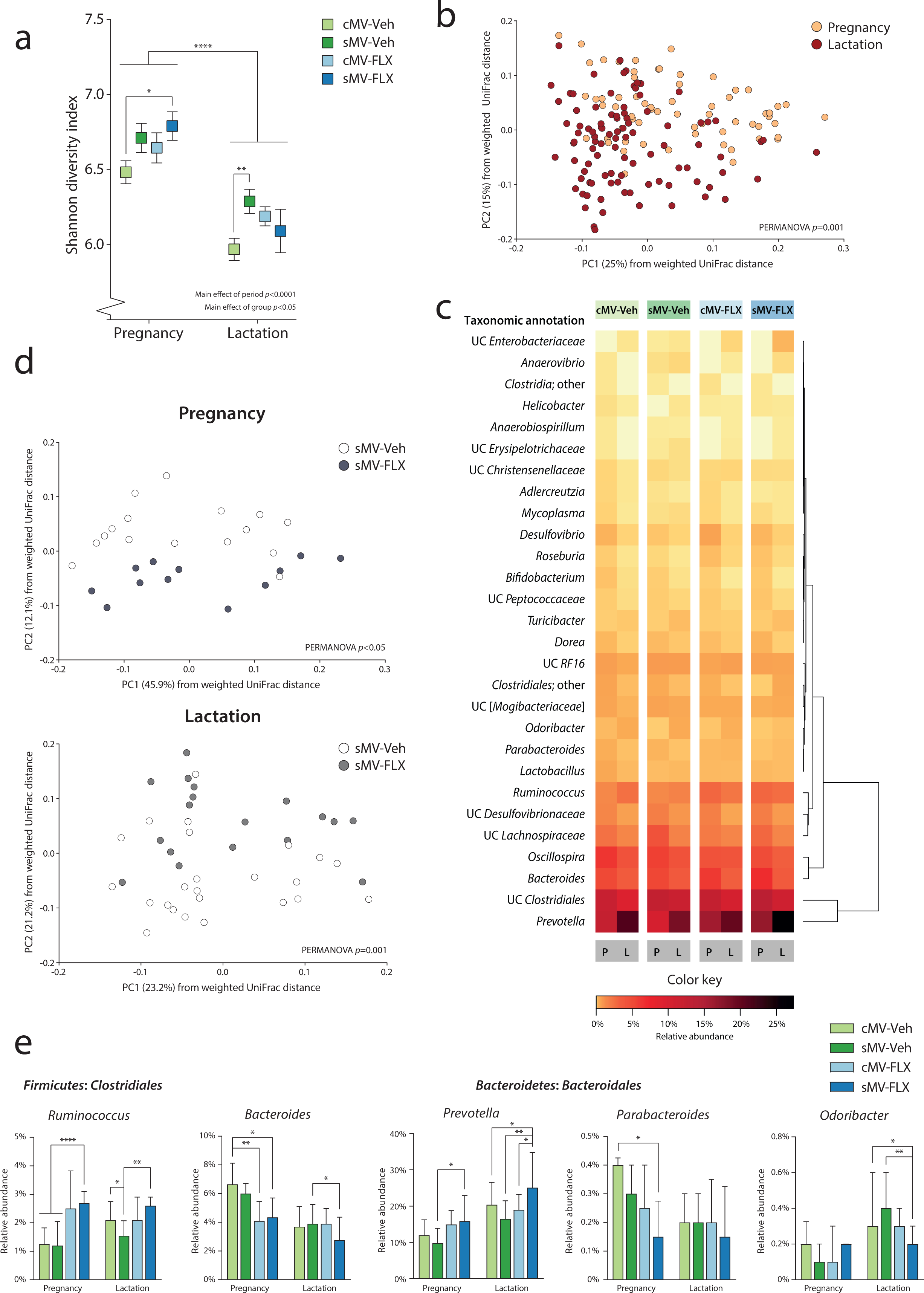
Fluoxetine treatment alters the microbiome during pregnancy and lactation in rats with a depressive-like phenotype. (a) Microbial alpha diversity. The Shannon diversity index was used as a measure of alpha diversity. A two-way ANOVA was performed with period (pregnancy vs lactation) and group as factors. Uncorrected Fisher’s LSD was used for group comparisons within periods. **(b) Structure of the microbial communities during pregnancy and lactation.** Communities were clustered using PCoA of the weighted UniFrac distance matrix. Each point corresponds to the microbial community of one sample that was collected during pregnancy or lactation. The percentage of variation explained by the PC is indicated on the axes. Colors correspond to period; all samples were grouped within pregnancy and lactation. **(c) Heatmap of relative abundances of Random Forests-identified OTUs distinguishing between pregnancy and lactation.** Random Forests was used to identify the microbial OTUs that had the highest predictive value for distinguishing between pregnancy and lactation in the cMV-Veh group. The heatmap depicts the relative abundance of these 28 OTUs during pregnancy and lactation for all groups, ordered by unsupervised clustering. **(d) The effect of FLX on the structure of the sMV microbiome during pregnancy and lactation.** Communities were clustered using PCoA of the weighted UniFrac distance matrix. Each point corresponds to the microbial community of one sample that was collected during pregnancy (upper graph) or lactation (lower graph). The percentage of variation explained by the PC is indicated on the axes. Colors correspond to group; sMV-Veh in white and sMV-FLX in grey. **(e) Relative abundance of selected Random Forests-identified OTUs.** Kruskal-Wallis tests were used, with subsequent group comparisons within periods (see Methods). N=12-22 samples/group from pregnancy (N=64 in total), N=18-33/group from lactation (N=96 in total).

In order to identify the bacterial signatures that underlie the observed shift in community structure between pregnancy and lactation, a machine-learning approach called Random Forests was used to analyze the data of the cMV-Veh group. This generated a model containing 28 Operational Taxonomic Units (OTUs) at the genus-level, discriminating between pregnancy and lactation. A large reorganization of the microbiome composition from gestation to lactation was revealed, consistent with structural remodeling (Figure 2c; Supplemental Figure 1a). Particularly within the *Bacteroidales* and *Clostridiales* orders, OTUs changed in their relative abundance from pregnancy to lactation (Figure 2c; Supplemental Figure 1a).

### Fluoxetine treatment alters the maternal fecal bacterial signatures of pregnancy and lactation in a rat model for depressive-like behavior

To test whether early life stress, FLX, and their combination alter the observed fecal microbial signatures of pregnancy and lactation as observed in the cMV-Veh group, we first compared the groups with respect to alpha- and beta diversity measures. We observed pre-conception differences between the sMV- and cMV group, with the sMV group showing higher alpha diversity than the cMV group (Supplemental Figure 2a, *p*<0.05), coinciding with differences in OTU relative abundance (Supplemental Figure 2b). During pregnancy and lactation there was a similar pattern for the sMV-Veh but also the cMV-FLX and sMV-FLX groups; the mean Shannon diversity of the fecal microbial communities of the treatment groups tended to be higher than that of the cMV-Veh group. However, this was only statistically significant for the sMV-FLX group during pregnancy and the sMV-Veh group during lactation (Figure 2a, pregnancy cMV-Veh vs sMV-Veh *p*=0.09; vs cMV-FLX *p*=0.25; vs sMV-FLX *p*<0.05 | lactation cMV-Veh vs sMV-Veh *p*<0.001; vs cMV-FLX *p*=0.06; vs sMV-FLX *p*=0.31). The treatment groups dissociated in terms of community structure, based on PCoA of the weighted UniFrac distances (Supplemental Figure 1b, PERMANOVA *p*=0.001). In particular, microbial community structure in the sMV-FLX group differed from the structure in the sMV-Veh group, both during pregnancy and lactation (Figure 1d, pregnancy sMV-Veh vs sMV-FLX PERMANOVA *p*<0.05 | lactation sMV-Veh vs sMV-FLX PERMANOVA *p*=0.001). This suggested that, in addition to the dynamic restructuring of the fecal microbiome across pregnancy and lactation, fluoxetine treatment modulated community structure of the fecal microbiota in the sMV females during these periods.

Next, the relative abundance of the 28 OTUs constituting the Random Forests-generated model discriminating between pregnancy and lactation in the cMV-Veh group was examined for the treatment groups. Globally, the pattern of OTUs distinguishing between pregnancy and lactation in the treatment groups was similar to the cMV-Veh group (Figure 2c). However, specific OTUs showed variations in their relative abundance within pregnancy and/or lactation (Figure 2e; Supplemental Figure 1c). The 2 OTUs with the largest contribution to the accuracy of the Random Forests model, *Bacteroides* and *Prevotella* from the *Bacteroidales* order, were sensitive to FLX treatment. Specifically, both the cMV-FLX and sMV-FLX had a lower relative abundance of *Bacteroides* than the cMV-Veh group during pregnancy, and the sMV-FLX group had a lower relative abundance of *Bacteroides* than the sMV-Veh group during lactation (Figure 2e, pregnancy cMV-Veh vs cMV-FLX *p*<0.01; vs sMV-FLX *p*<0.05 | lactation sMV-Veh vs sMV-FLX *p*<0.05). On the other hand, the sMV-FLX group had a higher relative abundance of *Prevotella* than the sMV-Veh group during pregnancy and relative to the cMV-Veh, sMV-Veh and cMV-FLX groups during lactation, reaching almost 25% of the total sample in lactation (Figure 2e, pregnancy sMV-Veh vs sMV-FLX *p*<0.05 | lactation cMV-Veh vs sMV-FLX *p*<0.05, sMV-Veh vs sMV-FLX *p*<0.01, cMV-FLX vs sMV-FLX *p*<0.05). Early life stress alone did not change the abundance of many OTUs relative to group that was control handled, although *Ruminococcus* was lower in the sMV-Veh group than in the cMV-Veh group during lactation (Figure 2e, lactation cMV-Veh vs sMV-Veh *p*<0.05). This effect was reversed by FLX, as the sMV-FLX had a higher relative abundance than cMV-Veh and sMV-Veh during pregnancy, and a higher relative abundance than sMV-Veh during lactation (Figure 2e, pregnancy cMV-Veh/ sMV-Veh vs sMV-FLX *p*<0.001 | lactation sMV-Veh vs sMV-FLX *p*<0.01).

Considering that OTUs within a community display co-occurrence and co-exclusion relationships, we performed a correlation analysis to look at these relationships within our samples [50]. A correlation matrix between the 28 selected OTUs revealed that *Prevotella* was negatively correlated with a cluster consisting mainly of *Clostridiales* (Supplemental Figure 1d). *Ruminoccus* was the exception within the *Clostridiales*, being positively correlated with *Prevotella* (Supplemental Figure 1d).

Together, these results confirm that pregnancy and lactation are distinct with regard to fecal microbiota diversity, structure, and composition. Early life stress increased alpha diversity during lactation, as was the case pre-conception. There were no profound effects of early life stress alone on either microbial community structure or on the relative abundance of OTUs resulting from our Random Forests model. However, FLX impacted community structure during pregnancy and lactation, as well as the relative abundance of a number of OTUs selected for their ability to distinguish between pregnancy and lactation. FLX in combination with early life stress had the largest effects on the maternal microbiome.

### Pregnancy and lactation have distinct fecal metabolic signatures

Bacterial communities metabolize and synthesize a number of metabolites that are essential for both dam and offspring. Shifts in bacterial community composition and predictive function may suggest alterations in the capacity of these communities to synthesize or metabolize these metabolites. We first performed PICRUSt predictive analysis on the OTU data to explore the hypothesis that the functional potential of pregnancy and lactation is different, since they have different microbial signatures. To determine which distinct predictive metabolic pathways discriminate between pregnancy and lactation, Random Forests analysis was performed on PICRUSt-generated data from the cMV-Veh group. Nine KEGG pathways were significantly changed across pregnancy and lactation, including the pathway of lysine biosynthesis; the pathway of alanine, aspartate, and glutamate metabolism; and the pathway of arginine and proline metabolism (Figure 3a; Supplemental Figure 3a). Overall, pathways related to amino acid synthesis and metabolism were among the metabolic pathways best able to characterize the difference in the predicted function of the maternal bacterial community during pregnancy vs lactation.

**Figure 3:**
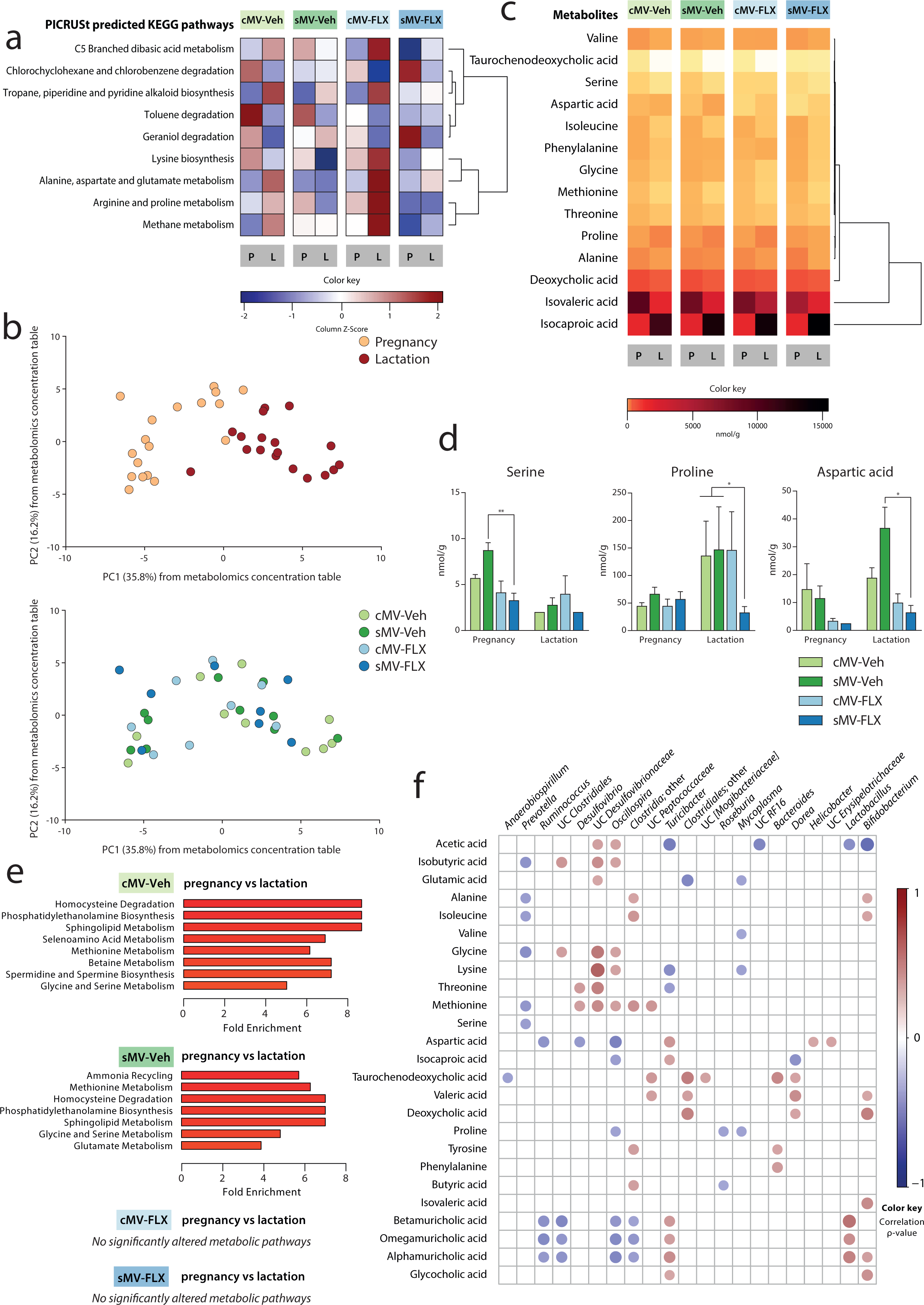
Fluoxetine alters fecal metabolite availability during pregnancy and lactation in rats with a depressive-like phenotype. **(a) Heatmap of Random Forests-identified PICRUSt-generated KEGG pathways distinguishing between pregnancy and lactation.** Z-scores are depicted of the abundance of the 9 identified pathways for all groups. The pathways are ordered by unsupervised clustering. **(b) Structure of the metabolomic composition during pregnancy and lactation.** Samples were clustered using PCA of the metabolomics concentration table. Each point corresponds to the metabolic capacity of one sample that was collected during pregnancy or lactation. The percentage of variation explained by the PC is indicated on the axes. Colors correspond to period in the upper graph, and to group in the lower graph. **(c) Heatmap of Random Forests-identified metabolite concentrations distinguishing between pregnancy and lactation.** Z-scores are depicted of the concentrations of the 14 identified metabolites for all groups. **(d) Concentrations of selected amino acids.** Kruskal-Wallis tests were used, with subsequent group comparisons within periods (see Methods). **(e) Metabolite Set Enrichment Analysis of pregnancy vs lactation per group**. Only the pathways that were significantly altered between pregnancy and lactation are shown here; the full list of metabolic pathways is shown in Supplemental Figure 3e. **(f) Correlation matrix of metabolites and OTUs throughout pregnancy and lactation.** Spearman’s rank correlations were determined between the 14 Random Forests-identified metabolites and the 28 Random Forests-identified OTUs. The circles represent the correlation between each metabolite and OTU. The size of the circle increases with decreasing *p*-value-, and the color of the circle corresponds to the ρ-value of the correlation (the strength of the correlation). Only significant (*p*<0.05) correlations are plotted, and only the OTUs with at least 1 significant correlation are shown here. The matrix is ordered by hierarchical clustering. For a: N=12-22 samples/group from pregnancy (N=64 in total), N=18-33/group from lactation (N=96 in total). For b-f: N=4-5/group from pregnancy (N=18 in total), N=4-5/group from lactation (N=18 in total).

To determine whether this altered predictive metabolic capacity of the microbiome was associated with actual metabolite availability, we measured the concentration of amino acids, bile acids, and short-chain fatty acids in a subset (N=4-5 per group in pregnancy and lactation) of fecal pellets. Corresponding to the microbiome results, principle component analysis (PCA) revealed a clear distinction in metabolic profile between samples obtained during pregnancy versus lactation (Figure 3b, upper graph).

To reveal the metabolic signatures of pregnancy and lactation, Random Forests analysis was performed on the metabolite data from the cMV-Veh group. Fourteen metabolites were included in the resulting model to distinguish between pregnancy and lactation (Supplemental Figure 3b). Most of them were amino acids, and 11 out of 14 metabolites were higher in concentration during pregnancy than during lactation (Figure 3c; Supplemental Figure 3c). For instance, the amino acids isoleucine and methionine had higher concentrations during pregnancy than during lactation, while proline concentrations were higher during lactation than during pregnancy (Figure 3c; Supplemental Figure 3c). The bile acids deoxycholic acid and taurochenodeoxycholic acid were both higher in concentration during pregnancy than during lactation (Figure 3c; Supplemental Figure 3c). Of the short-chain fatty acids included in the Random Forests model, isocaproic acid concentrations were very low during pregnancy, whereas isovaleric acid concentrations decreased during lactation (Figure 3c; Supplemental Figure 3c).

Next, we used Metabolite Set Enrichment Analysis (MSEA) to identify the fecal metabolite sets that were enriched during either pregnancy or lactation. In the cMV-Veh group, there were several pathways that significantly differed between these two periods, again containing amino acid-related pathways such as glycine and serine metabolism (Figure 3e; Supplemental Figure 3e).

### Fluoxetine treatment alters maternal fecal metabolite availability during pregnancy and lactation in a rat model for depressive-like behavior

To investigate whether early life stress, FLX and their combination affect the abovementioned fecal metabolic signatures of pregnancy and lactation as observed in the cMV-Veh group, we compared these groups with respect to predicted- and measured metabolic data. The effects of early life stress and FLX on the predictive pathways were mixed, with the sMV-FLX group having generally the lowest predicted abundance of these metabolic pathways (Figure 3a). Visual inspection of the PCA graph based on the metabolite concentration table revealed no clear effect of early life stress or FLX treatment (Figure 3b, lower graph).

Next, the concentrations of the 14 metabolites included in the Random Forests-generated model discriminating between pregnancy and lactation in the cMV-Veh group were examined for the treatment groups. Similar to the microbiome results, the overall pattern of metabolite availability changes between pregnancy and lactation was similar in all groups (Figure 3c). However, again, specific metabolites showed variations in their concentration within pregnancy and/or lactation (Figure 3d; Supplemental figure 3c). While there were no significant differences between the cMV-Veh and sMV-Veh group for any metabolites, the sMV-FLX group had low fecal concentrations of some amino acids (Figure 3d). Concentrations of serine were lower in the sMV-FLX group than in the sMV-Veh group during pregnancy, and for proline and aspartic acid this was the case during lactation (Figure 3d, serine pregnancy sMV-Veh vs sMV-FLX *p*<0.01, proline lactation cMV-Veh/ sMV-Veh vs sMV-FLX *p*<0.05, aspartic acid lactation sMV-Veh vs sMV-FLX *p*<0.05). From the metabolites that were not selected by the Random Forests model as distinguishing between pregnancy and lactation, the concentration of the bile acids betamuricholic acid and omegamuricholic acid were decreased during pregnancy in the sMV-FLX group compared to the cMV-Veh group and both the cMV-Veh- and the sMV-Veh group, respectively (Supplemental Figure 3d, betamuricholic acid pregnancy cMV-Veh vs cMV-FLX/sMV-FLX *p*<0.05, omegamuricholic acid pregnancy cMV-Veh/sMV-Veh vs sMV-FLX *p*<0.05). Similarly, the short-chain fatty acid isobutyric acid was decreased in the sMV-FLX group relative to the Ctrl- and the sMV-Veh group during lactation (Supplemental Figure 3d, cMV-Veh/sMV-Veh vs sMV-FLX *p*<0.05).

Then, we performed MSEA on the sMV-Veh, cMV-FLX and sMV-FLX data to determine which metabolites involved in specific metabolic processes were enriched in either pregnancy or lactation. The sMV-Veh group, like the cMV-Veh group, contained a number of metabolite sets that distinguished between pregnancy and lactation, such as methionine metabolism, and glycine and serine metabolism (Figure 3e; Supplemental Figure 3e). In contrast, in the cMV-FLX and sMV-FLX groups no metabolite sets were identified that differed significantly between pregnancy and lactation (Figure 3e; Supplemental Figure 3e).

Overall, the metabolomic results were consistent with the microbiome findings and the predictive functional analysis in discriminating between pregnancy and lactation and pointing towards amino acids as being central to this shift. Fluoxetine treatment lowered the fecal concentrations of several amino acids during these dynamic periods in our model for depressive-like symptoms. In addition, fluoxetine treatment abolished the differences in MSEA-identified pathways that distinguished the fecal metabolome during pregnancy from lactation as observed in the vehicle-treated groups.

### The relative abundances of fecal bacterial taxa correlate with metabolite concentrations

Finally, we generated a correlation matrix to further explore the link between the variation in the composition of the microbiome and variation in the availability of metabolites. A cluster consisting of OTUs from the *Clostridiales* (including *Oscillospira, Clostridia*;other and UC *Peptococcaceae*) and *Desulfovibrionales* (*Desulfovibrio* and UC *Desulfovibrionaceae*) orders was positively correlated with methionine, and some of these OTUs also to other amino acids such as alanine, isoleucine glycine, lycine, and threonine (Figure 3f). Showing almost the reverse pattern, the relative abundance of *Prevotella* was negatively correlated with availability of isobutyric acid and a collection of amino acids; alanine, isoleucine, glycine, methionine, and serine (Figure 3f). Associating to a different set of metabolites, the relative abundance of *Ruminococcus* was inversely correlated with concentrations of aspartic acid and some bile acids; betamuricholic acid, omegamuricholic acid, and alphamuricholic acid (Figure 3f). Overall, the OTUs that were most affected in relative abundance by FLX treatment (*Prevotella* and *Ruminococcus*, among others) are correlated with amino acid availability (serine, proline and aspartic acid, among others).

## Discussion

Our study shows that antidepressant treatment during pregnancy and lactation disrupts metabolite availability in a rat model for maternal depressive-like symptoms, possibly through alterations in bacterial community composition and function. In short, maternal vulnerability (MV) female rats were exposed to early life stress (sMV) in the form of maternal separation. In adulthood, sMV and control (cMV) females were treated throughout gestation and lactation with fluoxetine (FLX), a commonly prescribed selective serotonin reuptake inhibitor (SSRI), or vehicle (Veh). Fecal samples were collected during pregnancy and lactation, and 16s rRNA sequencing and targeted metabolomics were performed on these samples.

We first identified the distinct fecal microbial signatures of pregnancy and lactation in cMV-Veh females. We showed that the gut microbiome alpha diversity was relatively high during pregnancy, and dropped during lactation. In addition, principal coordinates analysis revealed a shift in the structure of the fecal microbiome from pregnancy to lactation. Using a machine-learning procedure called Random Forests we identified the set of microbial taxa that was overrepresented in either pregnancy or lactation. The resulting model, including 28 OTUs, characterized the shift in the maternal gut bacterial community between pregnancy and lactation. Twenty-three OTUs in the resulting model were higher in abundance during pregnancy than during lactation. In the literature there are several reports describing the changes in the maternal microbiome over the course of gestation in humans and rodents [23, 38, 51]. We extend these studies to define distinct microbial signatures associated with pregnancy and lactation.

Studies have shown that the mere presence of bacteria, but also the specific community composition in the gut greatly affects plasma and fecal metabolite availability [52, 53]. To begin to explore the hypothesis that, since pregnancy and lactation have different microbial signatures, their functional capacity is also different, we performed PICRUSt predictive analysis on the OTU data. Using Random Forests to determine the set of metabolic pathways that was best able to characterize the difference between pregnancy and lactation, we identified a set of 9 KEGG pathways, a large proportion of which was related to amino acid synthesis and metabolism. Gut microbes utilize metabolites as fuel, but also modify and produce short-chain fatty acids, bile acids and amino acids that are used by the host [54, 55]. We measured the concentrations of these metabolites in a subset of fecal samples. Random Forests analysis identified a set of 14 metabolites that best characterized the difference in fecal metabolite availability between pregnancy and lactation in cMV-Veh rats. Most of these metabolites – 11 out of 14 – were higher in concentration during pregnancy than during lactation, reflecting either greater intake or production of these metabolites during pregnancy, or more breakdown or transfer to the offspring during lactation. Maternal (bacterial) metabolites have been shown to transfer to the developing offspring through placental transfer and mother milk and play crucial roles in development [22, 56–58]. We postulate that the observed shift in metabolic profile from pregnancy to lactation is an adaptive one, aiding the pregnant and lactating female in providing nutrients and metabolites to the developing offspring.

We then hypothesized that early life stress and FLX alter the fecal microbial signatures of pregnancy and lactation, and that the combination of early life stress and FLX produces the most pronounced effect. An estimated 7-13% of women are diagnosed with a major depressive disorder in the perinatal period [28, 29]. Numerous studies have identified differences between the gut microbiome of depressed patients or depressive-like rodents, and healthy controls [13–15]. In this study, we used sMV females as a rat model with a depressive-like phenotype [32]. It has been shown repeatedly that maternal separation in rodents produces long-term effects not only on behavior, immune function and neurobiology, but also on the gut microbiome [59–61]. Correspondingly, our lab showed previously that maternal separation exacerbated the differences in the gut microbiome composition in young rats with a heterozygous or complete knockout of SERT compared to wildtypes [12]. Consistent with these results, we showed here that adult sMV females had a higher fecal alpha diversity than cMV females pre-conception, during pregnancy and during lactation [32]. However, maternal separation did not affect the structure and composition of the microbiome during pregnancy and lactation. It seems that the shift in maternal microbial community structure and composition from pregnancy to lactation is robust enough to withstand major perturbations by our early life stress protocol. In addition, no significant differences were found between the cMV-Veh and sMV-Veh groups in any metabolite concentration.

However, the SSRI fluoxetine was associated with differences in the maternal fecal microbiome and metabolic output in sMV rats. Around 1-10% of women use SSRI antidepressants during pregnancy [30, 31]. Medication has the capacity to profoundly alter the gut microbiome composition [62, 63]. Indeed, there are several *in vitro* studies suggesting that SSRIs possess antimicrobial properties [16–19]. Moreover, it was shown recently that psychotropic drugs, including fluoxetine, are capable of altering the composition of the gut microbiome in rats [20]. Our results here confirm that SSRIs have the potential to modulate the fecal microbiome during pregnancy and lactation in rats. However, we only find significant effects of FLX on maternal gut microbial community dynamics in sMV-, and not in cMV females. Principle coordinates analysis showed an effect of FLX on the sMV bacterial community structure. Similarly, when we compared the relative abundance of the Random Forests-identified taxa between the treatments, most effects were seen in the sMV-FLX group relative to the sMV-Veh group. For instance, fecal relative abundances of *Prevotella* and *Ruminococcus* were higher in the sMV-FLX group than in the sMV-Veh group, at the cost of *Bacteroides*.

Thus, treatment with fluoxetine during pregnancy and lactation modulated the microbiome almost exclusively in the sMV animals, and not significantly in the cMV animals. There are several possible explanations for this. It might be that the sMV microbiome, belonging to a depressive-like host, is more sensitive to disruptions than the cMV microbiome, especially during the dynamic perinatal period. It might also be that both early life stress and FLX are “hits” that can shift the balance in the gut microbial community, and that under the current experimental conditions these “hits” are not large enough to be observed individually, but only when they are combined (a double-hit model). At the moment, there is a paucity of studies into the effect of SSRI on the microbiome from clinical studies on depression or animal studies mimicking this condition. For instance, it is unclear whether any effects SSRIs might have on the microbiome are part of their therapeutic potential or should rather be seen as side effects.

Then, we were interested in whether the effects of FLX on the maternal fecal microbiome would be reflected in altered metabolite concentrations. Indeed, FLX treatment suppressed the fecal availability of some amino acids in sMV females. Specifically, we found levels of serine to be lower in sMV-FLX animals compared to sMV-Veh animals during pregnancy, and proline and aspartic acid to be lower in the sMV-FLX group compared to the sMV-Veh group during lactation. Similarly, a study using the unpredictable mild stress paradigm showed that in stressed or depressive-like rats, serine plasma levels are higher than in controls, while fluoxetine treatment lowered serine plasma levels compared to controls [64]. That study, however, did not include an experimental group receiving both stress and fluoxetine [64]. Bacteria are able to produce amino acids from nonspecific nitrogen sources [65]. Amino acids play essential roles in both homeostatic physiology in adults as well as development and survival of the fetus, and are involved in cell growth and differentiation, protein synthesis, hormone secretion and lactation [66, 67]. During early pregnancy, maternal amino acids are only second to glucose in the amount of substrate that crosses the placenta to support the growth of the developing fetus [68].

To further examine the metabolic pathways that were characteristic of either pregnancy or lactation based on fecal metabolite concentrations and the effect of early life stress and FLX on these pathways, metabolite set enrichment analysis was performed. Both the cMV-Veh and the sMV-Veh group showed a clear signature of pregnancy vs lactation, reflected in a number of metabolic pathways (including amino acid-related pathways) that were enriched in either period. Both the cMV-FLX and the sMV-FLX group showed no such significantly enriched pathways, adding to the emerging picture that FLX disrupts the fecal metabolic signatures of pregnancy and lactation. Future studies should address whether this results in altered metabolite transfer to the offspring and what the possible consequences are for offspring development.

Finally, to link the 16S rRNA sequencing data with the metabolomic data, we generated correlation matrices both between the 28 OTUs selected by Random Forests analysis, as well as between the relative abundance of these OTUs and metabolite concentrations. We found that the relative abundance of *Prevotella*, enhanced in the sMV-FLX group, was negatively correlated with a cluster of *Clostridiales* and *Desulfovibrionales* genera. Interestingly, these same *Clostridiales* and *Desulfovibrionales* OTUs were positively associated with availability of amino acids. Previous literature describes that some of the most prevalent amino acid fermenting microbes indeed belong to the *Clostridiales* and *Proteobacteria* taxa [69]. The associations we report here yield a working hypothesis that FLX – at least in an animal model for depressive-like behavior – tips the balance toward higher abundance *Prevotella* and lower abundance of the aforementioned cluster of amino acid-fermenting OTUs, which leads to lower levels of amino acids. Further mechanistic studies are needed to test this hypothesis.

This study has several limitations. As with any 16S rRNA marker gene sequencing approach, our analysis was constrained to the genus level. In addition, we are aware that the microbiome and its functional capacity varies across locations in the gastrointestinal tract whereas we only used fecal samples [70]. However, our study design enabled us to sample longitudinally and to reduce the number of animals needed to complete the study. Another issue is the observed mortality in about one-fourth of our SSRI-treated females. This may be the result of the SERT^+/-^ genotype of the MV females. Indeed, humans carrying the short allele of the SERT gene polymorphism are more likely to suffer from side effects from SSRIs than controls without a short allele [71]. In future studies, wildtype SERT^+/+^ will have to be included in order to establish whether this is indeed a genotype X drug effect. None of the females included in the current study showed any signs of toxicity as a result of FLX. For future studies we also suggest taking plasma samples, and possibly mother’s milk, to confirm whether changes in fecal metabolite concentrations correspond to systemic availability of these metabolites.

The benefits of performing microbiome research on laboratory rodents as opposed to humans deserve mentioning here. For example, diet composition could be kept stable throughout the study, whereas diet composition changes considerably from pre-conception to pregnancy in humans and even short-term changes in diet are known to affect the gut microbiome [72, 73]. In addition, we are able to study the independent effect of SSRI use in rodents, as well as its interactions with a depressive-like phenotype. Human SSRI-users always have underlying psychopathologies that are difficult to control for [74]. However, the biggest asset of the current study is our ability to link changes to the microbiome and predicted functional capacity of the bacterial community to changes in metabolite availability. Especially in the context of pregnancy and lactation, any changes to maternal microbial metabolite availability might have far-reaching consequences and are therefore important to assess in addition to the microbiome itself.

Our results add to the growing body of research linking serotonin signaling to the gut microbiome. Maternal depression and SSRI use during pregnancy are both associated with detrimental developmental outcomes in offspring. Clinical and animal studies have identified adverse effects such as lower birth weight, delayed motor development, and increased anxiety [75]. Future studies will assess whether and to what extent the maternal gut microbiome and metabolome might mediate these effects. Microbial molecules have been shown to transfer from mother to offspring during gestation and through lactation to support the development of the brain and the immune system, emphasizing their importance for offspring development [21, 22, 76, 77]. If the dampening effect of SSRI treatment on fecal amino acid concentrations in a rodent model of depressive-like symptoms can be replicated in future studies, and if these concentrations correlate with maternal and fetal plasma levels and to phenotypic outcomes in offspring, dietary supplementation of amino acids alongside SSRI treatment during pregnancy and lactation might be an important direction for future research.

## Conclusions

We showed here that the fecal microbial and metabolic signatures of pregnancy and lactation in MV rats are robust and are largely unaltered by early life stress with maternal separation. However, fluoxetine treatment modulated key aspects of maternal microbial community dynamics and metabolite output in the sMV females, including changes in the relative abundance of OTUs and associated decreases in amino acid availability. This suggests that antidepressant use during pregnancy and lactation may lead to changes in the microbiome. The current study adds to the emerging awareness that the microbiome is plastic and vulnerable to environmental and pharmacological insults, especially during dynamic periods such as pregnancy and lactation. We speculate that alterations to the maternal microbiome by SSRI treatment, and the associated decreased availability of amino acids, reflect a compromised ability to supply nutrients to the developing offspring.

## List of abbreviations

cMV: maternal vulnerability females exposed to control handling
FLX: fluoxetine(-treated)
IQR: interquartile range
KEGG: Kyoto Encyclopedia of Genes and Genomes
L: lactation
MSEA: Metabolite Set Enrichment Analysis
OTU: operational taxonomic unit
P: pregnancy
PCA: principal component analysis
PC: principal component/coordinate
PCoA: principle coordinates analysis
PICRUSt: Phylogenetic Investigation of Communities by Reconstruction of Unobserved States
SCFA: short-chain fatty acids
sMV: maternal vulnerability females exposed to early life stress
UC: unclassified
Veh: vehicle-treated.

## Declarations

### Ethics approval and consent to participate

The animal study was approved by the Institutional Animal Care and Use Committee at the University of Groningen (no. DEC-6936A), and was conducted in accordance with the institutional guidelines and the Law on Animal Experiments.

### Consent for publication

Not applicable.

### Authors’ contributions

ASR, EJ, TLB and JDAO designed the study. ASR and DJH performed the animal experiment and collected the samples. ASR and EJ performed the wet lab work and bioinformatics analysis. ASR drafted the manuscript. All authors contributed to critical revision of the manuscript.

## Acknowledgements

We thank Judith Swart, Wanda Douwenga and Christa Reitzema-Klein for their assistance with the early life stress procedure, drug administration and sample collection.

---

**Supplemental Figure S1:**
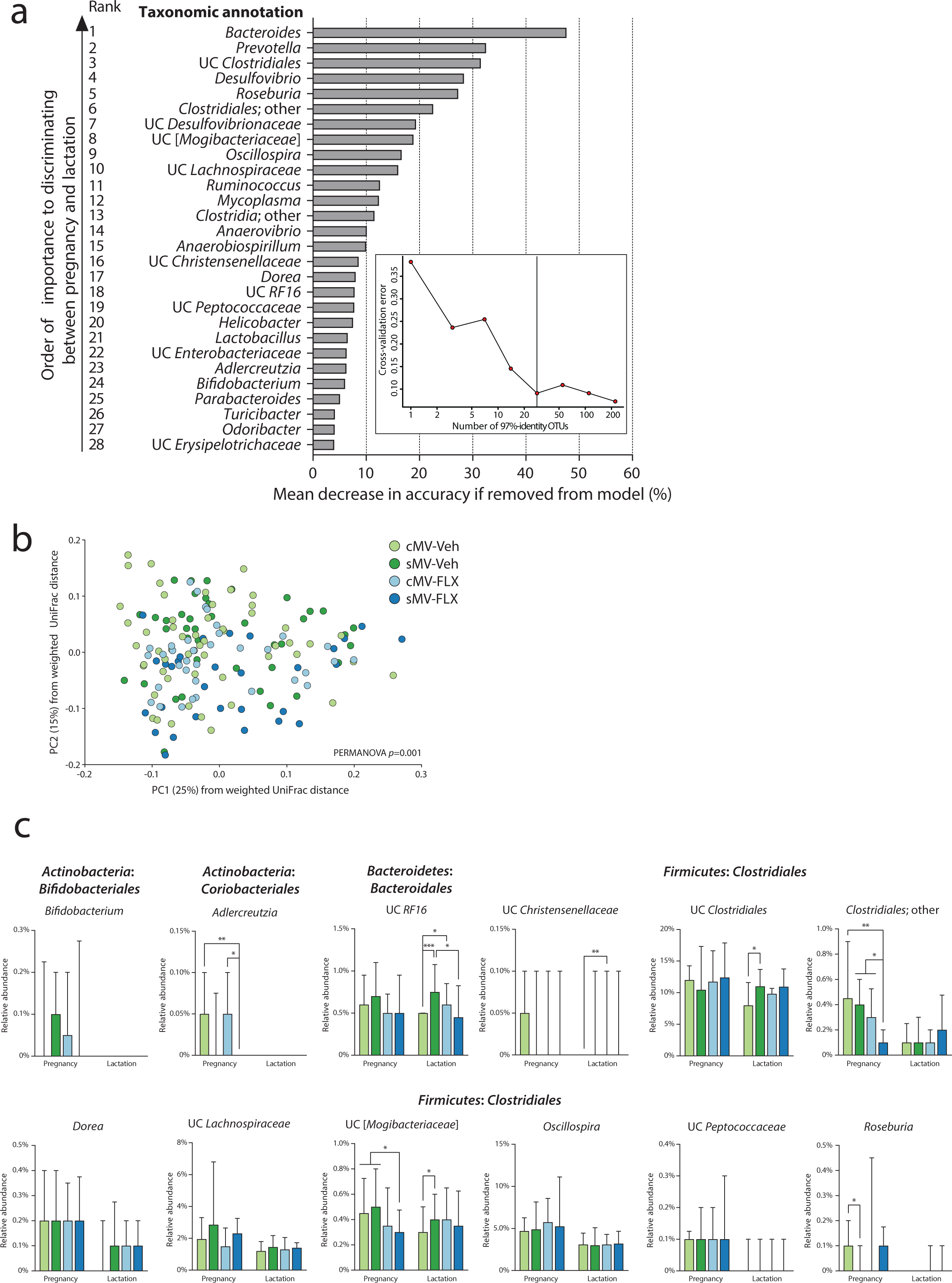
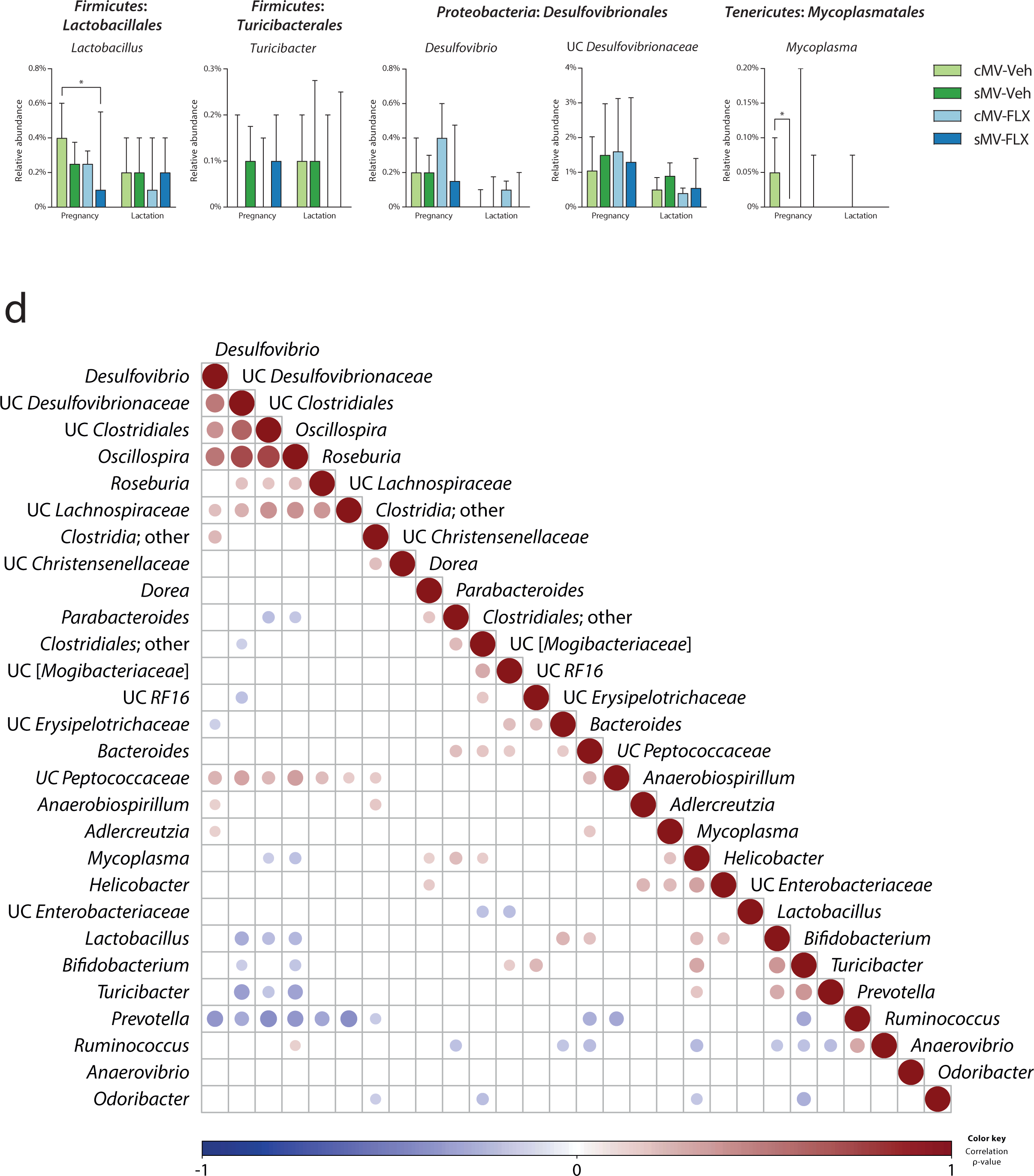
The effect of early life stress and fluoxetine on the microbiome during pregnancy and lactation. **(a) Random Forests model to determine which OTUs were best able to distinguish between pregnancy and lactation in the cMV-Veh group.** The inset figure shows increasing numbers of 97%-identity OTUs on the x-axis plotted against the cross-validation error of the model on the y-axis. It was determined that the optimal model would include the top 28 OTUs, which are shown in the bigger bar graph. From top to bottom, the OTUs are shown in descending order, corresponding to their importance to discriminating between pregnancy and lactation. The length of the bars represents the mean decrease in accuracy of the model if this OTU would be removed from the model. **(b) Structure of the microbial communities during pregnancy and lactation.** Communities were clustered using PCoA of the weighted UniFrac distance matrix. Each point corresponds to the microbial community of one sample that was collected during pregnancy or lactation. The percentage of variation explained by the PC is indicated on the axes. Colors correspond to group (containing samples from both pregnancy and lactation). **(c) Bar graphs showing relative abundance of selected Random Forests-identified OTUs.** A Kruskal-Wallis test was used, with subsequent group comparisons within periods (see Methods). Only OTUs with a median relative abundance > 0 in at least 1 of the groups are shown here. **(d) Correlation matrix of OTUs throughout pregnancy and lactation.** Spearman’s rank correlations were determined between the 28 Random Forests-identified OTUs. The circles represent the correlation between two given OTUs, with the size of the circle increasing with decreasing *p*-value of the correlation, and the color of the circle corresponding to the ρ-value of the correlation (the strength of the correlation). Only significant (*p*<0.05) correlations are plotted. The OTUs are ordered by hierarchical clustering. N=12-22 samples/group from pregnancy (N=64 in total), N=18-33/group from lactation (N=96 in total).

**Supplemental Figure S2:**
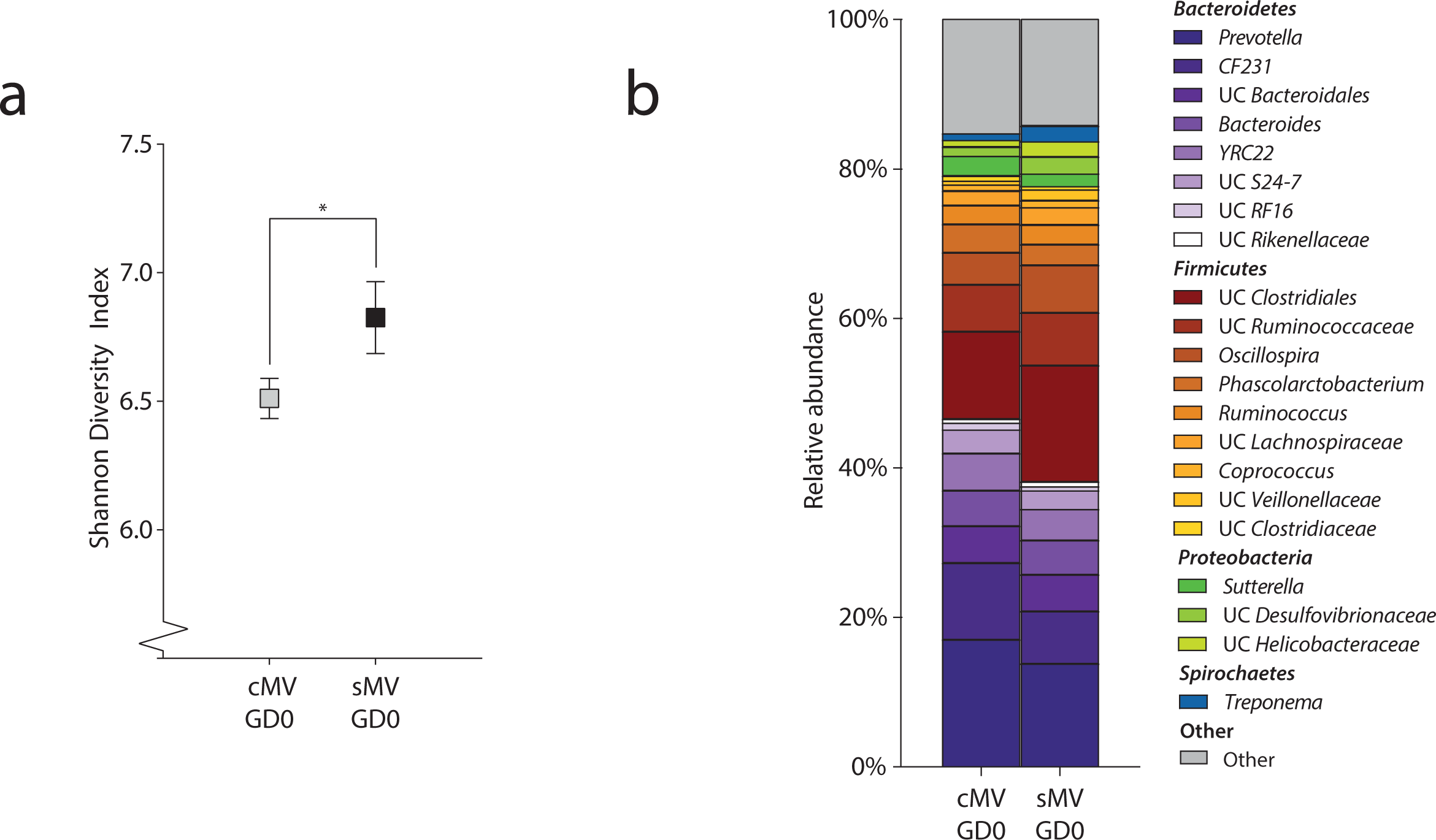
The effect of early life stress on the gut microbiome in adult MV female rats. **a) Pre-conception alpha diversity.** A t-test was used to test whether adult cMV and sMV females differed in terms of fecal microbial Shannon Diversity. **(b) Pre-conception microbiome composition.** A stack plot of the 20 most abundant OTUs was used to compare the fecal microbiome composition of cMV and sMV females. N=18 for cMV; N=14 for sMV.

**Supplemental Figure S3:**
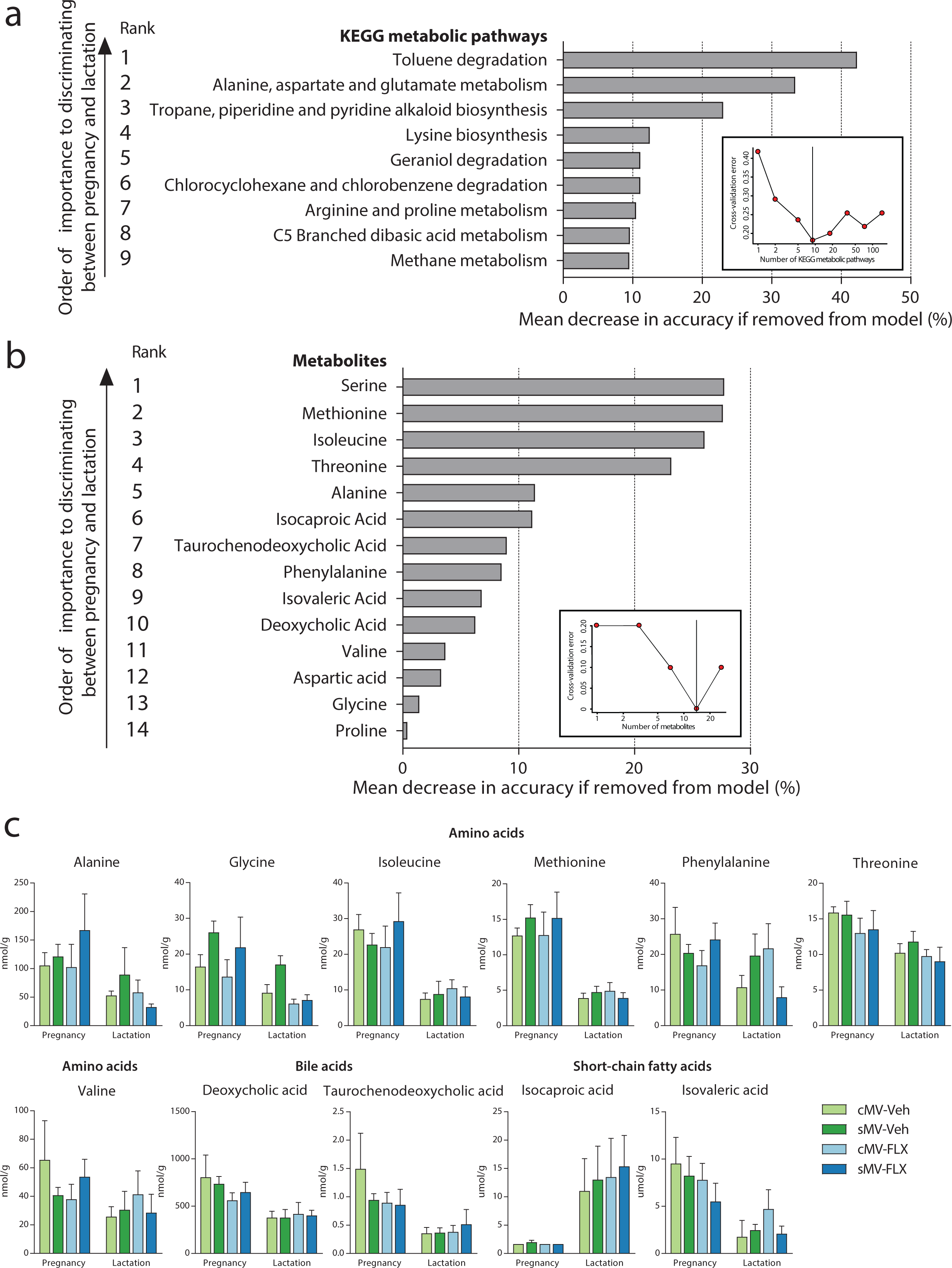
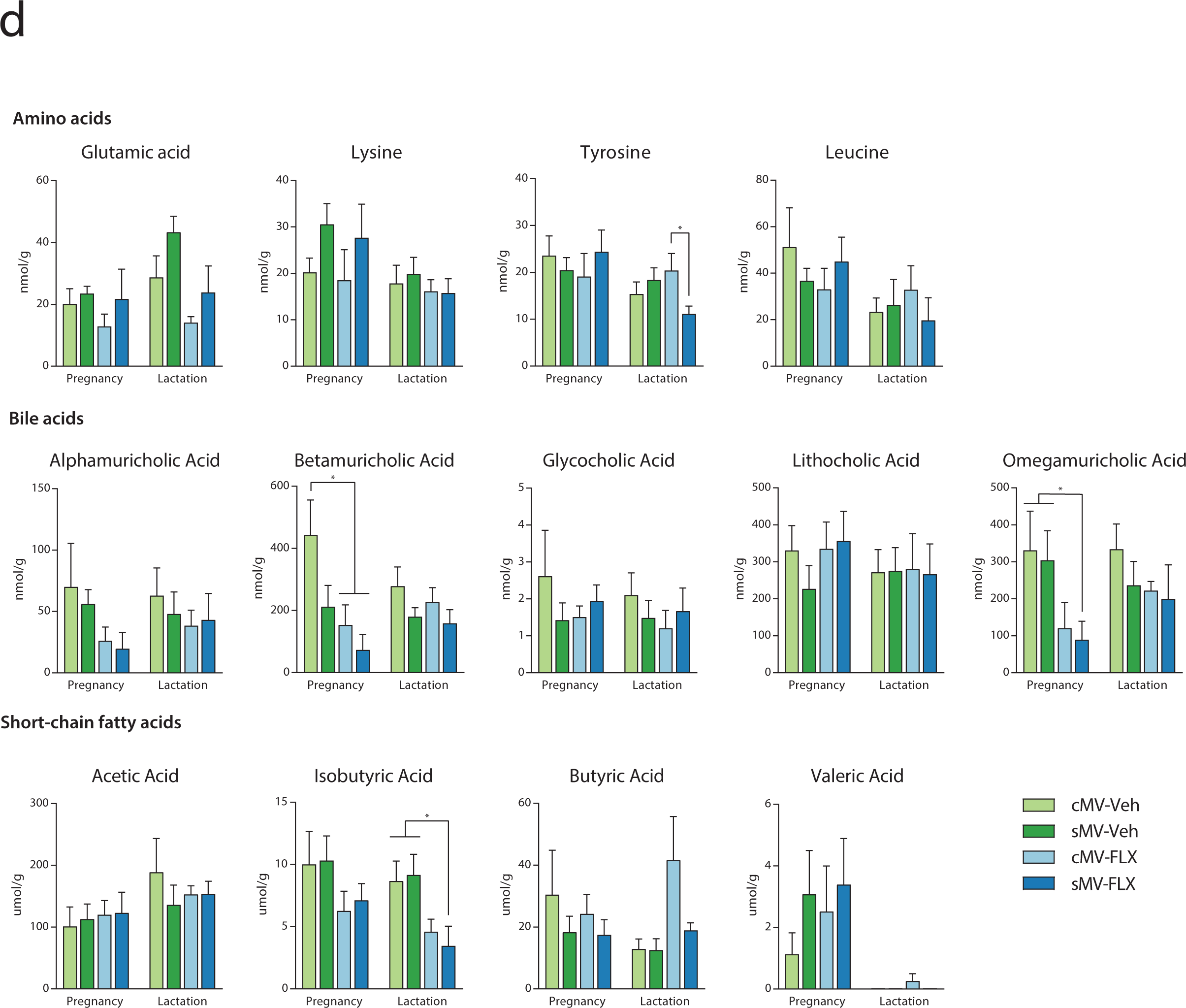
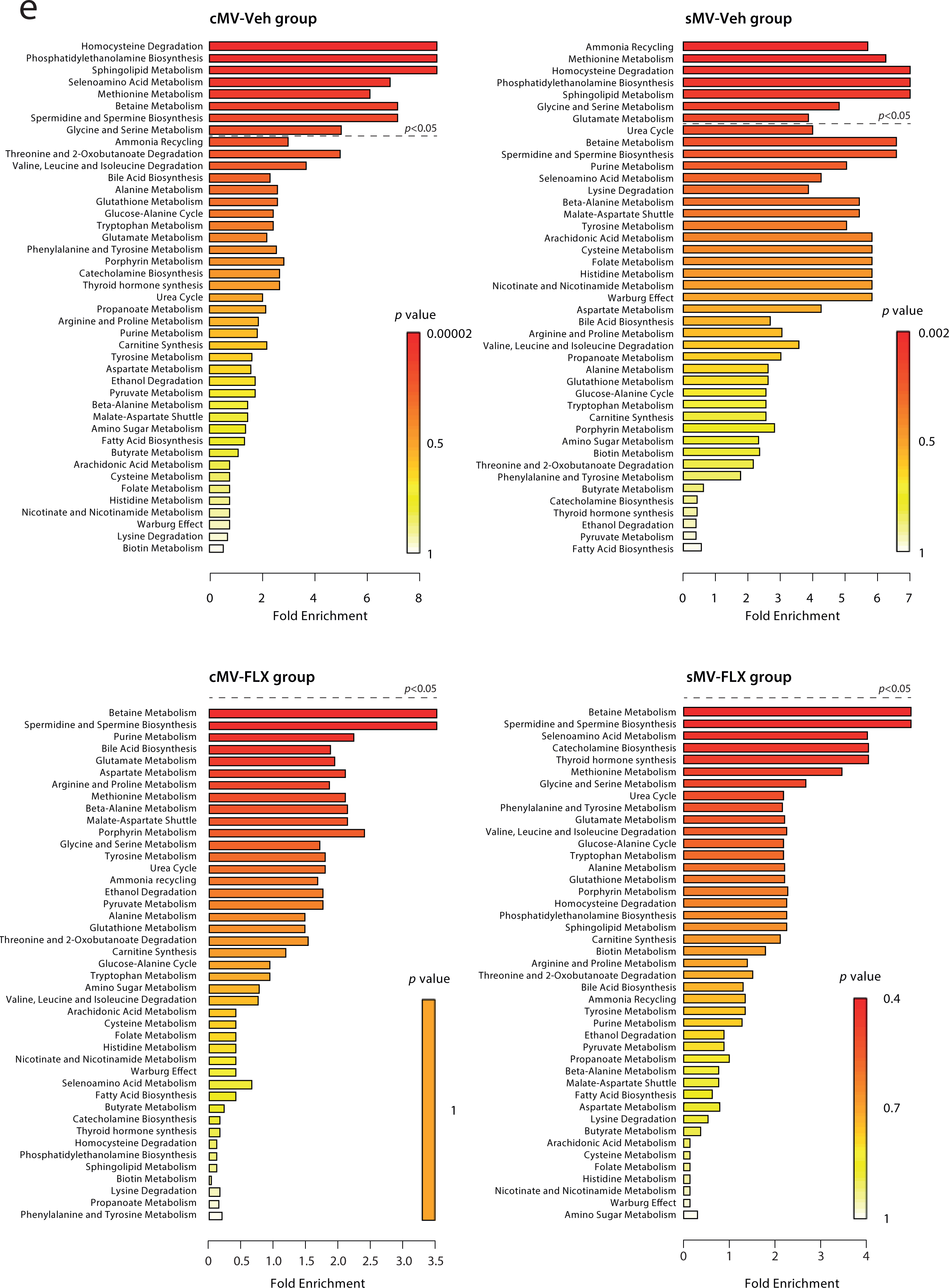
The effect of early life stress and fluoxetine on metabolite availability during pregnancy and lactation. **(a) Random Forests model to determine which predicted KEGG metabolic pathways were best able to distinguish between pregnancy and lactation in the cMV-Veh group.** The inset figure shows increasing numbers of KEGG metabolic pathways on the x-axis plotted against the cross-validation error of the model on the y-axis. It was determined that the optimal model would include the top 9 KEGG metabolic pathways. The length of the bars represents the mean decrease in accuracy of the model if this KEGG pathway would be removed from the model. **(b) Random Forests model to determine which metabolites were best able to distinguish between pregnancy and lactation in the cMV-Veh group.** The smaller figure shows increasing numbers metabolites on the x-axis plotted against the cross-validation error of the model on the y-axis. It was determined that the optimal model would include the top 14 metabolites, which are shown in the bigger bar graph. The length of the bars represents the mean decrease in accuracy of the model if this metabolite would be removed from the model. **(c) Bar graphs showing concentrations of Random Forests-identified metabolites.** A Kruskal-Wallis test was used, with subsequent group comparisons within periods (see Methods). **(d) Bar graphs showing concentrations of the non-Random Forests-identified metabolites.** A Kruskal-Wallis test was used, with subsequent group comparisons within periods (see Methods). **(e) Metabolite Set Enrichment Analysis of pregnancy vs lactation per group**. MSEA was used to identify the metabolic pathways distinguishing pregnancy and lactation based on the metabolomics concentration table for each group. For a: N=22 cMV-Veh samples from pregnancy and N=33 from lactation. For b: N=5 cMV-Veh samples from pregnancy and N=5 from lactation. For c: N=4-5/group from pregnancy and N=4-5/group from lactation (N=36 in total).

## References

1. Coppen A. The biochemistry of affective disorders. Br J Psychiatry. 1967;113:1237–64. doi:10.1192/bjp.113.504.1237.

2. Fakhoury M. Revisiting the serotonin hypothesis: Implications for major depressive disorders. Mol Neurobiol. 2016;53:2778–86. doi:10.1007/s12035-015-9152-z.

3. Claassen V, Davies JE, Hertting G, Placheta P. Fluvoxamine, a Specific 5-Hydroxytryptamine Uptake Inhibitor. Br J Pharmacol. 1977;60:505–16.

4. Gershon MD, Tack J. The Serotonin Signaling System: From Basic Understanding To Drug Development for Functional GI Disorders. Gastroenterology. 2007;132:397–414. doi:10.1053/j.gastro.2006.11.002.

5. Sjögren K, Engdahl C, Henning P, Lerner UH, Tremaroli V, Lagerquist MK, et al. The gut microbiota regulates bone mass in mice. J Bone Miner Res. 2012;27:1357–67. doi:10.1002/jbmr.1588.

6. Ge X, Ding C, Zhao W, Xu L, Tian H, Gong J, et al. Antibiotics - induced depletion of mice microbiota induces changes in host serotonin biosynthesis and intestinal motility. J Transl Med. 2017;:1–9.

7. Hata T, Asano Y, Yoshihara K, Kimura-todani T, Miyata N, Zhang X, et al. Regulation of gut luminal serotonin by commensal microbiota in mice. 2017;:1–14.

8. Reigstad CS, Salmonson CE, Rainey JF, Szurszewski JH, Linden DR, Sonnenburg JL, et al. Gut microbes promote colonic serotonin production through an effect of short-chain fatty acids on enterochromaffin cells. FASEB J. 2015;29:1395–403. doi:10.1096/fj.14-259598.

9. Yano JM, Yu K, Donaldson GP, Shastri GG, Ann P, Ma L, et al. Indigenous bacteria from the gut microbiota regulate host serotonin biosynthesis. Cell. 2015;161:264–76. doi:10.1016/j.cell.2015.02.047.

10. Spohn SN, Mawe GM. Non-conventional features of peripheral serotonin signalling — the gut and beyond. Nat Publ Gr. 2017. doi:10.1038/nrgastro.2017.51.

11. O’Mahony SM, Clarke G, Borre YE, Dinan TG, Cryan JF. Serotonin, tryptophan metabolism and the brain-gut-microbiome axis. Behav Brain Res. 2015;277:32–48. doi:10.1016/j.bbr.2014.07.027.

12. El Aidy S, Ramsteijn AS, Dini-Andreote F, van Eijk R, Houwing DJ, Salles JF, et al. Serotonin transporter genotype modulates the gut microbiota composition in young rats, an effect augmented by early life stress. Front Cell Neurosci. 2017;11 August:1–12. doi:10.3389/fncel.2017.00222.

13. Naseribafrouei A, Hestad K, Avershina E, Sekelja M, Linløkken A, Wilson R, et al. Correlation between the human fecal microbiota and depression. Neurogastroenterol Motil. 2014;26:1155–62.

14. Jiang H, Ling Z, Zhang Y, Mao H, Ma Z, Yin Y, et al. Altered fecal microbiota composition in patients with major depressive disorder. Brain Behav Immun. 2015;48:186–94. doi:10.1016/j.bbi.2015.03.016.

15. Park AJ, Collins J, Blennerhassett PA, Ghia JE, Verdu EF, Bercik P, et al. Altered colonic function and microbiota profile in a mouse model of chronic depression. Neurogastroenterol Motil. 2013;25.

16. Muñoz-Bellido JL, Muñoz-Criado S, García-Rodríguez JA. In-vitro activity of psychiatric drugs against Corynebacterium urealyticum (Corynebacterium group D2). J Antimicrob Chemother. 1996;37:1005–9.

17. Bohnert JA, Szymaniak-Vits M, Schuster S, Kern W V. Efflux inhibition by selective serotonin reuptake inhibitors in Escherichia coli. J Antimicrob Chemother. 2011;66:2057–60.

18. Macedo D, Filho Ajmc, Soares de Sousa CN, Quevedo J, Barichello T, Júnior HVN, et al. Antidepressants, antimicrobials or both? Gut microbiota dysbiosis in depression and possible implications of the antimicrobial effects of antidepressant drugs for antidepressant effectiveness. J Affect Disord. 2017;208 September 2016:22–32. doi:10.1016/j.jad.2016.09.012.

19. Evrensel A, Ceylan ME. Gut-Microbiota-Brain Axis and Depression. In: Kim Y-K, editor. Understanding Depression. Singapore: Springer Singapore; 2018. p. 197–207. doi:10.1007/978-981-10-6580-4_17.

20. Cussotto S, Strain CR, Fouhy F, Strain RG, Peterson VL, Clarke G, et al. Differential effects of psychotropic drugs on microbiome composition and gastrointestinal function. Psychopharmacology (Berl). 2018;:1–15. doi:10.1007/s00213-018-5006-5.

21. Braniste V, Al-Asmakh M, Kowal C, Anuar F, Abbaspour A, Toth M, et al. The gut microbiota influences blood-brain barrier permeability in mice. Sci Transl Med. 2014;6:263ra158–263ra158. doi:10.1126/scitranslmed.3009759.

22. Gomez de Aguero M, Ganal-Vonarburg SC, Fuhrer T, Rupp S, Uchimura Y, Li H, et al. The maternal microbiota drives early postnatal innate immune development. Science (80-). 2016;351:1296–302. doi:10.1126/science.aad2571.

23. Jašarević E, Howard CD, Misic AM, Beiting DP, Bale TL. Stress during pregnancy alters temporal and spatial dynamics of the maternal and offspring microbiome in a sex-specific manner. Sci Rep. 2017;7 March:44182. doi:10.1038/srep44182.

24. Gohir W, Whelan FJ, Surette MG, Moore C, Jonathan D, Sloboda DM, et al. Pregnancy-related changes in the maternal gut microbiota are dependent upon the mother’s periconceptional diet. Gut Microbes. 2015;0976 January 2016:310–20.

25. Khan I, Azhar EI, Abbas AT, Kumosani T, Barbour EK, Raoult D, et al. Metagenomic analysis of antibiotic-induced changes in gut microbiota in a pregnant rat model. Front Pharmacol. 2016;7 APR:1–11.

26. Mandal S, Godfrey KM, McDonald D, Treuren W V., Bjørnholt J V., Midtvedt T, et al. Fat and vitamin intakes during pregnancy have stronger relations with a proinflammatory maternal microbiota than does carbohydrate intake. Microbiome. 2016;4:1–11. doi:10.1186/s40168-016-0200-3.

27. Tochitani S, Ikeno T, Ito T, Sakurai A, Yamauchi T, Matsuzaki H. Administration of non-Absorbable antibiotics to pregnant mice to perturb the maternal gut microbiota is associated with alterations in offspring behavior. PLoS One. 2016;11:1–22.

28. Gavin NI, Gaynes BN, Lorh KN, Meltzer-Brody S, Gartlehner G, Swinson T. Perinatal depression: A systematic review of prevalence and incidence. Obstet Gynecol. 2005;106:1071–83.

29. Woody CA, Ferrari AJ, Siskind DJ, Whiteford HA, Harris MG. A systematic review and meta-regression of the prevalence and incidence of perinatal depression. J Affect Disord. 2017;219 May:86–92. doi:10.1016/j.jad.2017.05.003.

30. El Marroun H, Jaddoe VW V., Hudziak JJ, Roza SJ, Steegers EAP, Hofman A, et al. Maternal use of selective serotonin reuptake inhibitors, fetal growth, and risk of adverse birth outcomes. Arch Gen Psychiatry. 2012;69:706–14. doi:10.1001/archgenpsychiatry.2011.2333.

31. Cooper WO, Willy ME, Pont SJ, Ray WA. Increasing use of antidepressants in pregnancy. Am J Obstet Gynecol. 2007;196:544.e1-5.

32. Houwing DJ, Ramsteijn AS, Riemersma IW, Olivier JDA. Maternal separation induces anhedonia in female heterozygous serotonin transporter knockout rats. Manuscr Submitt Publ. 2018.

33. Smits BMG, Mudde JB, van de Belt J, Verheul M, Olivier J, Homberg J, et al. Generation of gene knockouts and mutant models in the laboratory rat by ENU-driven target-selected mutagenesis. Pharmacogenet Genomics. 2006;16:159–69.

34. Homberg JR, Olivier JDA, Smits BMG, Mul JD, Mudde J, Verheul M, et al. Characterization of the serotonin transporter knockout rat: A selective change in the functioning of the serotonergic system. Neuroscience. 2007;146:1662–76.

35. Haberstick BC, Smolen A, Williams RB, Bishop GD, Foshee VA, Thornberry TP, et al. Population Frequencies of the Triallelic 5HTTLPR in Six Ethnicially Diverse Samples from North America, Southeast Asia, and Africa. Calcif Tissue Int. 2015;96:255–61.

36. Caspi A, Sugden K, Moffitt TE, Taylor A, Craig IW, Harrington H, et al. Influence of life stress on depression: Moderation by a polymorphism in the 5-HTT gene. Science. 2003;301:386–9.

37. Houwing DJ, Buwalda B, van der Zee EA, de Boer SF, Olivier JDA. The Serotonin Transporter and Early Life Stress: Translational Perspectives. Front Cell Neurosci. 2017;11 April:1–16. doi:10.3389/fncel.2017.00117.

38. DiGiulio DB, Callahan BJ, McMurdie PJ, Costello EK, Lyell DJ, Robaczewska A, et al. Temporal and spatial variation of the human microbiota during pregnancy. Proc Natl Acad Sci. 2015;112:11060–5. doi:10.1073/pnas.1502875112.

39. Kozich JJ, Westcott SL, Baxter NT, Highlander SK, Schloss PD. Development of a dual-index sequencing strategy and curation pipeline for analyzing amplicon sequence data on the miseq illumina sequencing platform. Appl Environ Microbiol. 2013;79:5112–20.

40. Schloss PD, Westcott SL, Ryabin T, Hall JR, Hartmann M, Hollister EB, et al. Introducing mothur: Open-source, platform-independent, community-supported software for describing and comparing microbial communities. Appl Environ Microbiol. 2009;75:7537–41.

41. Caporaso JG, Kuczynski J, Stombaugh J, Bittinger K, Bushman FD, Costello EK, et al. QIIME allows analysis of high-throughput community sequencing data. Nat Methods. 2010;7:335–6. doi:10.1038/nmeth.f.303.

42. R: A language and environment for computing. 2017. https://www.r-project.org/.

43. Liaw a, Wiener M. Classification and Regression by randomForest. R news. 2002;2 December:18–22.

44. Subramanian S, Huq S, Yatsunenko T, Haque R, Mahfuz M, Alam MA, et al. Persistent gut microbiota immaturity in malnourished Bangladeshi children. Nature. 2014;510:417–21. doi:10.1038/nature13421.

45. Warnes GR, Bolker B, Bonebakker, Lodewijk Gentleman R, Huber W, Liaw A, Lumley T, et al. gplots: Various R programming tools for plotting data. 2016. https://cran.r-project.org/package=gplots.

46. Langille MGI, Zaneveld J, Caporaso JG, McDonald D, Knights D, Reyes JA, et al. Predictive functional profiling of microbial communities using 16S rRNA marker gene sequences. Nat Biotechnol. 2013;31:814–21. doi:10.1038/nbt.2676.

47. Xia J, Sinelnikov I V., Han B, Wishart DS. MetaboAnalyst 3.0-making metabolomics more meaningful. Nucleic Acids Res. 2015;43:W251–7.

48. Subramanian A, Tamayo P, Mootha VK, Mukherjee S, Ebert BL, Gillette MA, et al. Gene set enrichment analysis: A knowledge-based approach for interpreting genome-wide expression profiles. Proc Natl Acad Sci. 2005;102:15545–50. doi:10.1073/pnas.0506580102.

49. Anderson MJ, Connell SD, Gillanders BM, Diebel CE, Blom WM, Saunders JE, et al. A new method for non-parametric multivariate analysis of variance. Austral Ecol. 2001;26:32–46. doi:10.1111/j.1442-9993.2001.01070.pp.x.

50. Faust K, Sathirapongsasuti JF, Izard J, Segata N, Gevers D, Raes J, et al. Microbial co-occurrence relationships in the Human Microbiome. PLoS Comput Biol. 2012;8.

51. Koren O, Goodrich JK, Cullender TC, Spor A, Laitinen K, Kling Bäckhed H, et al. Host remodeling of the gut microbiome and metabolic changes during pregnancy. Cell. 2012;150:470–80.

52. Wikoff WR, Anfora AT, Liu J, Schultz PG, Lesley S a, Peters EC, et al. Metabolomics analysis reveals large effects of gut microflora on mammalian blood metabolites. Proc Natl Acad Sci U S A. 2009;106:3698–703. doi:10.1073/pnas.0812874106.

53. Marcobal A, Kashyap PC, Nelson TA, Aronov PA, Donia MS, Spormann A, et al. A metabolomic view of how the human gut microbiota impacts the host metabolome using humanized and gnotobiotic mice. ISME J. 2013;7:1933–43. doi:10.1038/ismej.2013.89.

54. Lin R, Liu W, Piao M, Zhu H. A review of the relationship between the gut microbiota and amino acid metabolism. Amino Acids. 2017;49:2083–90.

55. Nieuwdorp M, Gilijamse PW, Pai N, Kaplan LM. Role of the microbiome in energy regulation and metabolism. Gastroenterology. 2014;146:1525–33. doi:10.1053/j.gastro.2014.02.008.

56. van de Pavert SA, Ferreira M, Domingues RG, Ribeiro H, Molenaar R, Moreira-Santos L, et al. Maternal retinoids control type 3 innate lymphoid cells and set the offspring immunity. Nature. 2014;508:123–7. doi:10.1038/nature13158.

57. Thorburn AN, McKenzie CI, Shen S, Stanley D, MacIa L, Mason LJ, et al. Evidence that asthma is a developmental origin disease influenced by maternal diet and bacterial metabolites. Nat Commun. 2015;6.

58. Scholtens DM, Bain JR, Reisetter AC, Muehlbauer MJ, Nodzenski M, Stevens RD, et al. Metabolic networks and metabolites underlie associations between maternal glucose during pregnancy and newborn size at birth. Diabetes. 2016;65:2039–50.

59. Bailey MT, Coe CL. Maternal separation disrupts the integrity of the intestinal microflora in infant rhesus monkeys. Dev Psychobiol. 1999;35:146–55.

60. De Palma G, Blennerhassett P, Lu J, Deng Y, Park a. J, Green W, et al. Microbiota and host determinants of behavioural phenotype in maternally separated mice. Nat Commun. 2015;6 August:7735. doi:10.1038/ncomms8735.

61. Pusceddu MM, El Aidy S, Crispie F, O’Sullivan O, Cotter P, Stanton C, et al. N-3 polyunsaturated fatty acids (PUFAs) reverse the impact of early-life stress on the gut microbiota. PLoS One. 2015;10:1–13.

62. Devkota S. Prescription drugs obscure microbiome analyses. Science (80-). 2016;351:452–3.

63. Falony G, Joossens M, Vieira-Silva S, Wang J, Darzi Y, Faust K, et al. Population-level analysis of gut microbiome variation. Science (80-). 2016;352:560–4.

64. Zhao L, Xiong Z, Lu X, Zheng S, Wang F, Ge L, et al. Metabonomic evaluation of chronic unpredictable mild stress-induced changes in rats by intervention of fluoxetine by HILIC-UHPLC/MS. PLoS One. 2015;10:1–14.

65. Metges CC. Contribution of microbial amino acids to amino acid homeostasis of the host. J Nutr. 2000;130:1857S–64S. doi:10.1093/jn/130.7.1857S.

66. Wu G, Wu Z, Dai Z, Yang Y, Wang W, Liu C, et al. Dietary requirements of “nutritionally non-essential amino acids” by animals and humans. Amino Acids. 2013;44:1107–13.

67. Lin G, Wang X, Wu G, Feng C, Zhou H, Li D, et al. Improving amino acid nutrition to prevent intrauterine growth restriction in mammals. Amino Acids. 2014;46:1605–23.

68. Herrera E. Metabolic adaptations in pregnancy and their implications for the availability of substrates to the fetus. Eur J Clin Nutr. 2000;54:S47–51.

69. Dai Z-L, Wu G, Zhu W-Y. Amino acid metabolism in intestinal bacteria: links between gut ecology and host health. Front Biosci (Landmark Ed. 2011;16 September:1768–86. http://www.ncbi.nlm.nih.gov/pubmed/21196263.

70. Li D, Chen H, Mao B, Yang Q, Zhao J, Gu Z, et al. Microbial Biogeography and Core Microbiota of the Rat Digestive Tract. Sci Rep. 2017;8 March:1–16. doi:10.1038/srep45840.

71. Luddington NS, Mandadapu A, Husk M, El-Mallakh RS. Clinical Implications of Genetic Variation in the Serotonin Transporter Promoter Region. Prim Care Companion J Clin Psychiatry. 2009;11:93–102. doi:10.4088/PCC.08r00656.

72. Hillier SE, Olander EK. Women’s dietary changes before and during pregnancy: A systematic review. Midwifery. 2017;49 June 2016:19–31.

73. David LA, Maurice CF, Carmody RN, Gootenberg DB, Button JE, Wolfe BE, et al. Diet rapidly and reproducibly alters the human gut microbiome. Nature. 2014;505:559–63. doi:10.1038/nature12820.

74. Olivier JDA, Åkerud H, Sundström Poromaa I. Antenatal depression and antidepressants during pregnancy: Unraveling the complex interactions for the offspring. Eur J Pharmacol. 2015;753:257–62. doi:10.1016/j.ejphar.2014.07.049.

75. Olivier JDA, Akerud H, Kaihola H, Pawluski JL, Skalkidou A, Högberg U, et al. The effects of maternal depression and maternal selective serotonin reuptake inhibitor exposure on offspring. Front Cell Neurosci. 2013;7 May:73. doi:10.3389/fncel.2013.00073.

76. Thion MS, Low D, Silvin A, Chen J, Grisel P, Schulte-Schrepping J, et al. Microbiome influences prenatal and adult microglia in a sex-specific manner. Cell. 2017;:1–17. doi:10.1016/j.cell.2017.11.042.

77. Jašarević E, Howard CD, Morrison K, Misic A, Weinkopff T, Scott P, et al. The maternal vaginal microbiome partially mediates the effects of prenatal stress on offspring gut and hypothalamus. Nat Neurosci. 2018. doi:10.1038/s41593-018-0182-5.

